# Adapterama II: Universal amplicon sequencing on Illumina platforms (TaggiMatrix)

**DOI:** 10.1101/619544

**Authors:** Travis C. Glenn, Todd W. Pierson, Natalia J. Bayona-Vásquez, Troy J. Kieran, Sandra L. Hoffberg, Jesse C. Thomas, Daniel E. Lefever, John W. Finger, Bei Gao, Xiaoming Bian, Swarnali Louha, Ramya T. Kolli, Kerin Bentley, Julie Rushmore, Kelvin Wong, Timothy I. Shaw, Michael J. Rothrock, Anna M. McKee, Tai L. Guo, Rodney Mauricio, Marirosa Molina, Brian S. Cummings, Lawrence H. Lash, Kun Lu, Gregory S. Gilbert, Stephen P. Hubbell, Brant C. Faircloth

## Abstract

Next-generation sequencing (NGS) of amplicons is used in a wide variety of contexts. Most NGS amplicon sequencing remains overly expensive and inflexible, with library preparation strategies relying upon the fusion of locus-specific primers to full-length adapter sequences with a single identifying sequence or ligating adapters onto PCR products. In *Adapterama I*, we presented universal stubs and primers to produce thousands of unique index combinations and a modifiable system for incorporating them into Illumina libraries. Here, we describe multiple ways to use the *Adapterama* system and other approaches for amplicon sequencing on Illumina instruments. In the variant we use most frequently for large-scale projects, we fuse partial adapter sequences (TruSeq or Nextera) onto the 5’ end of locus-specific PCR primers with variable-length tag sequences between the adapter and locus-specific sequences. These fusion primers can be used combinatorially to amplify samples within a 96-well plate (eight forward primers + 12 reverse primers yield 8 × 12 = 96 combinations), and the resulting amplicons can be pooled. The initial PCR products then serve as template for a second round of PCR with dual-indexed iTru or iNext primers (also used combinatorially) to make full-length libraries. The resulting quadruple-indexed amplicons have diversity at most base positions and can be pooled with any standard Illumina library for sequencing. The number of sequencing reads from the amplicon pools can be adjusted, facilitating deep sequencing when required or reducing sequencing costs per sample to an economically trivial amount when deep coverage is not needed. We demonstrate the utility and versatility of our approaches with results from six projects using different implementations of our protocols. Thus, we show that these methods facilitate amplicon library construction for Illumina instruments at reduced cost with increased flexibility. A simple web page to design fusion primers compatible with iTru primers is available at: http://baddna.uga.edu/tools-taggi.html. A fast and easy to use program to demultiplex amplicon pools with internal indexes is available at: https://github.com/lefeverde/Mr_Demuxy.

## Introduction

Next-generation DNA sequencing (NGS) has facilitated a wide variety of benefits in the life sciences (Ansorg, 2009; Goodwin, McPherson & McCombie, 2016), and NGS instruments have an ever-growing capacity to generate more reads per run. Substantial progress has been made in developing new, lower-cost instruments, but much less progress has been made in reducing the cost of sequencing runs (cf., Glenn, 2011 vs. Glenn, 2016). Thus, the large number of reads from a typical NGS run comes with a relatively large buy-in cost but yields an extremely low cost per read. Frustratingly, within every NGS platform, the lowest-cost sequencing kits have the highest costs per read (Glenn, 2011; 2016). This creates a fundamental challenge: how do we efficiently create and pool large numbers of samples so that we can divide the cost of high capacity NGS sequencing runs among many samples, thereby reducing the cost per sample?

It is well known that identifying DNA sequences (commonly called indexes, tags, or barcodes; we use the term indexes throughout) can be incorporated during sample preparation for NGS (i.e., library construction) so that multiple samples can be pooled prior to NGS, thereby allowing the sequencing costs to be divided among the samples (see Faircloth & Glenn, 2012 and references therein). When sufficient unique identifying indexes are available, many samples, including samples from multiple projects, can be pooled and sequenced on higher throughput platforms which minimizes costs for all samples in the pool.

In many potential NGS applications, the number of desired reads per sample is limited, so the cost of preparing samples for NGS sequencing becomes the largest component of the overall cost of collecting sequence data. Thus, it is desirable to increase the number of low-cost library preparation methods available. As the cost of library construction is reduced, projects requiring fewer DNA sequences per sample become effective to conduct using NGS (e.g., if sample preparation plus sequencing for NGS is < sample preparation plus sequencing on capillary machines, then it is economical to switch).

### Previous NGS amplicon library preparation methods

Amplicon library preparations for NGS have been integrating indexes for more than a decade (e.g., Binladen et al., 2007; Craig et al., 2008). Early NGS strategies consisted of conducting individual PCRs targeting different DNA regions from one sample and then pooling them together. Then, full-length adapters would be ligated to each sample pool, providing sample-specific identifiers. This approach has the advantage of being economical regarding amplicon production, primer cost, and pooling of amplicons prior to adapter ligation, as well as being ecumenical because the resulting amplicons can be ligated to adapters for any sequencing platform. The downside of this first approach is that adapters must be ligated to the amplicons, which is time-consuming, expensive, and error-prone, and which can introduce errors into the resulting sequences. To avoid ligation of adapters to amplicons, most NGS amplicon sequencing strategies have subsequently relied upon the fusion of locus-specific primers to full-length adapter sequences and the addition of identical indexes to both 5’ and 3’ ends (e.g., Roche fusion primers; Binladen et al., 2007; Bentley et al., 2009; Bybee et al., 2011; Cronn et al., 2012; Shokralla et al., 2014). These strategies often use the whole sequencing run for amplicons only. Illumina platforms have traditionally struggled to sequence amplicons because: 1) the platform requires a diversity of bases at each base position (Mitra et al., 2015), which is easily achieved in genomic libraries but not in amplicon libraries; and 2) read-lengths are limited, making the complete sequencing of long amplicons challenging or impossible.

Several alternatives have been proposed to resolve the first issue (i.e., low base-diversity). Users have typically added a genomic library (e.g., the PhiX control library supplied by Illumina) to amplicon library pools to create the base-diversity needed, but this method wastes sequencing reads on non-target (PhiX) library. Second, to solve the issue of limited read-length, described above, custom sequencing primers can be used in place of the Read1 and/or Read2 sequencing primer(s) (Caporaso et al., 2011). This method allows for longer effective read-lengths by removing the read-length wasted by sequencing the primers used for amplification (e.g., 16S primer sequences), but it can be very expensive to optimize custom sequencing primers, costing hundreds of dollars for each attempt. Another alternative is to use the amplicons as template for shotgun library preparations, most often using Nextera library preparation kits (Illumina 2018a). A fourth method is to add heterogeneity spacers to the indexes in the form of one, two, three (etc.) bases before the index sequence (e.g., Cruaud et al., 2017), but because amplicons can contain repeats longer than the heterogeneity spacers, it is still possible to have regions of no diversity. Thus, all of the proposed solutions have specific limitations, and none are particularly economical for sequencing standard PCR products from a wide range of samples, as is typical in molecular ecology projects.

### NGS amplicon needs

In general, NGS has been widely adopted to sequence complex amplicon pools where cloning would have been used previously (e.g., 16S from bacterial communities or viruses within individuals). Such amplicon pools may have extensive or no length variation. Amplicons for single loci from haploid or diploid organisms (with no length variation between alleles) are typically still sequenced via capillary electrophoresis at a cost of about $5 USD per read. In contrast to the high cost of individual sequencing reads via capillary instruments, >50,000 paired-end reads can be obtained for $5 USD on the Illumina MiSeq. Unfortunately, MiSeq runs come in units of ~$2,000 USD for reads that total a length similar to that of capillary sequencing (Glenn, 2016; paired-end (PE) 300 reads). Thus, it would be desirable to have processes that allow users to: 1) pool samples from multiple projects on a single MiSeq run and divide costs proportionately, and 2) prepare templates (i.e., construct libraries) at costs less than or similar to those of traditional capillary sequencing.

Characteristics of an ideal system include: 1) use of universal Illumina sequencing primers; 2) minimizing total sample costs, ideally to be below standard capillary/Sanger sequencing; 3) minimizing time and equipment needed for library preparations; 4) minimizing buy-in (start-up) costs; 5) eliminating error-prone steps, such as adapter ligation, 6) maximizing the number of samples (e.g., ≥ thousands) that can be identified in a pool of samples run simultaneously, 7) maximizing the range of amplicons that can be added to other pools (e.g., from <1% to >90%), and 8) creating a very large universe of sample identifiers (e.g., ≥ millions) so that identifiers would not need to be shared among samples, studies, or researchers, even when coming through large sequencing centers.

Single-locus amplicon sequencing represents one extreme example of the needs identified above. In some scenarios, researchers may only be sequencing a single short, homogeneous amplicon where ≥ 20x coverage is excessive. The cost of sequencing reagents for only 20 reads of 600 bases on an Illumina MiSeq using version 3 chemistry, which generates ~20 million reads, is <$0.01 USD (i.e., 1 millionth of the run). It is impractical to amass 1 million amplicon samples for a single run. However, a small volume of dozens or hundreds of samples can be easily added into a MiSeq run with other samples/pools that need the remaining of reads. By paying the proportional sequencing costs for such projects, the cost of constructing libraries and conducting quality control on the libraries becomes the largest component of the total cost of collecting NGS data. Having the ability to combine libraries of many different kinds of samples, each with their own identification indexes, is critical to the feasibility of this strategy. We have developed, and describe below, a system to meet most of the design characteristics enumerated above.

In this paper, we focus on library preparation methods for amplicons. We introduce TaggiMatrix, which is an amplicon library preparation protocol that is built upon methods developed in *Adapterama I* (Glenn et al., 2019). This general method can be optimized for various criteria, including the minimization of library preparation cost and reduction of PCR bias. Briefly, by tagging both the forward and reverse locus-specific primers with different, variable-length index sequences, and also by including indexes in the iTru or iNext primers, we create quadruple-indexed libraries with high base-diversity, enabling the use of highly combinatorial strategies to index, pool, and sequence many samples on Illumina instruments.

## Materials & Methods

### Methodological objectives

Our goal was to develop a protocol that would help to overcome the challenges of amplicon library preparation and fulfill the characteristics of an ideal system enumerated above. We extend the work of Faircloth & Glenn (2012) and Glenn et al. (2019) to achieve these goals.

### Methodological approach

Illumina libraries require four sequences (P5 + Read1 and P7 + Read2; Fig. 1), and can accommodate internal index sequences on each end, (i.e., P5 + i5 index + Read1 and P7 + i7 index + Read2; Fig. 1; Illumina Sequencing Dual-Indexed Libraries on the HiSeq System User Guide; Glenn et al., 2019). The Read1 and Read2 sequences can be of two types—TruSeq or Nextera—. Just as in *Adapterama I* (Glenn et al., 2019), we have designed systems for both.

**Figure 1.**
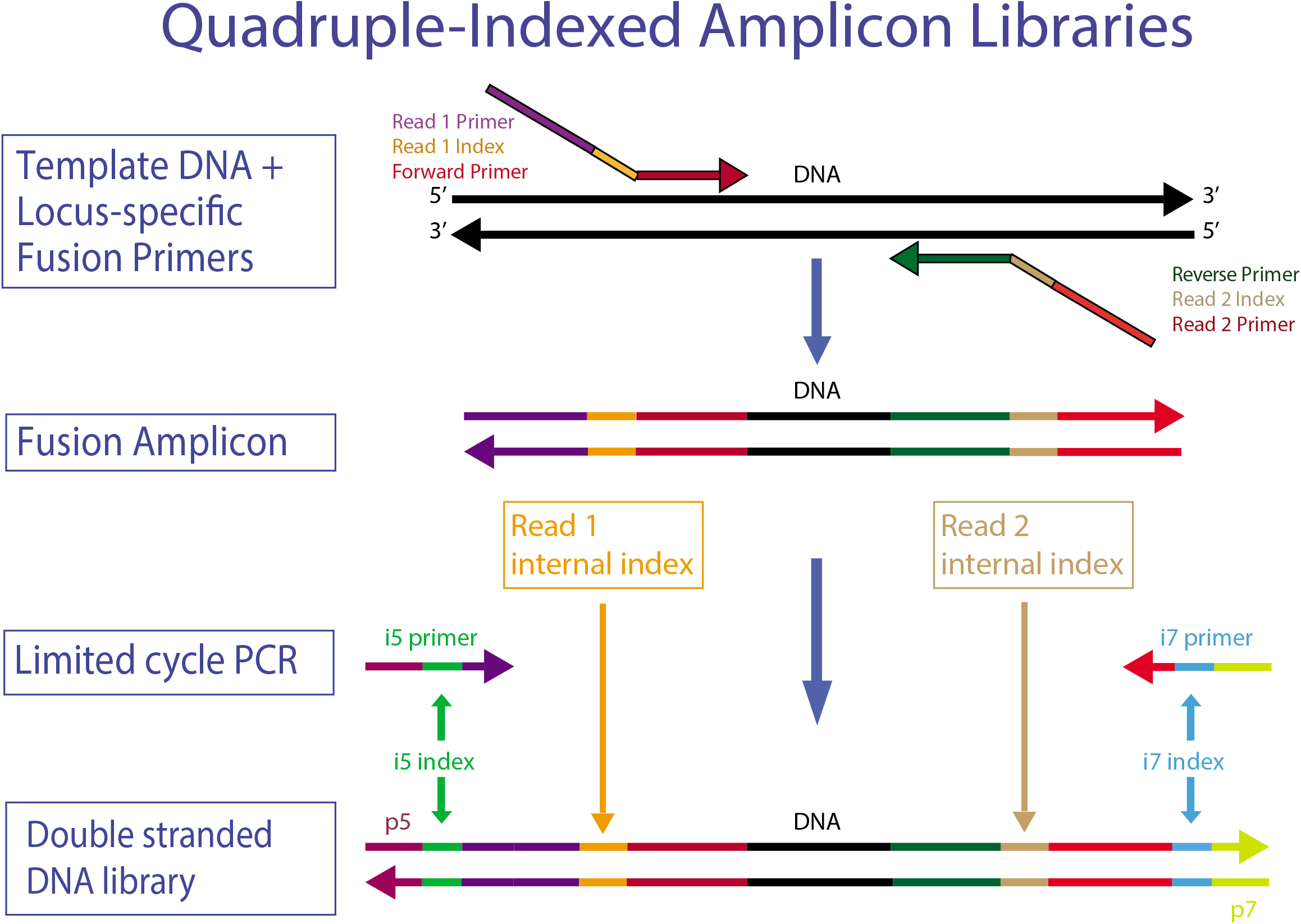
High throughput workflow to create and multiplex TaggiMatrix libraries. The components of the quadrupled-indexed amplicon Libraries. A specific DNA region is amplified using fusion and tagged locus-specific primers, also known as “indexed fusion primers”, to produce a fusion amplicon. Then iTru adapters are ligated using Y-yolk adapters or incorporated using limited cycle PCR with i5 and i7 indexed primers to make the complete double stranded DNA library. Internal indexes and outer i5/i7 indexes are represented as well as the set of primers used.

Our overall approach is to make amplicons with fusion primers (Fig. 2) that can use iTru or iNext primers described in *Adapterama I* (Glenn et al., 2019) to make full-length Illumina libraries (Fig. 3a; Figs. S1 and S2). The resulting libraries always contain dual-indexes in the standard indexing positions and may optionally contain additional internal indexes (Figs. 1–3; Table 1; Illumina, 2018b). These indexes are recovered through the four standard separate sequencing reactions generated by Illumina instruments when doing paired-end sequencing (Fig. 3b).

**Figure 2.**
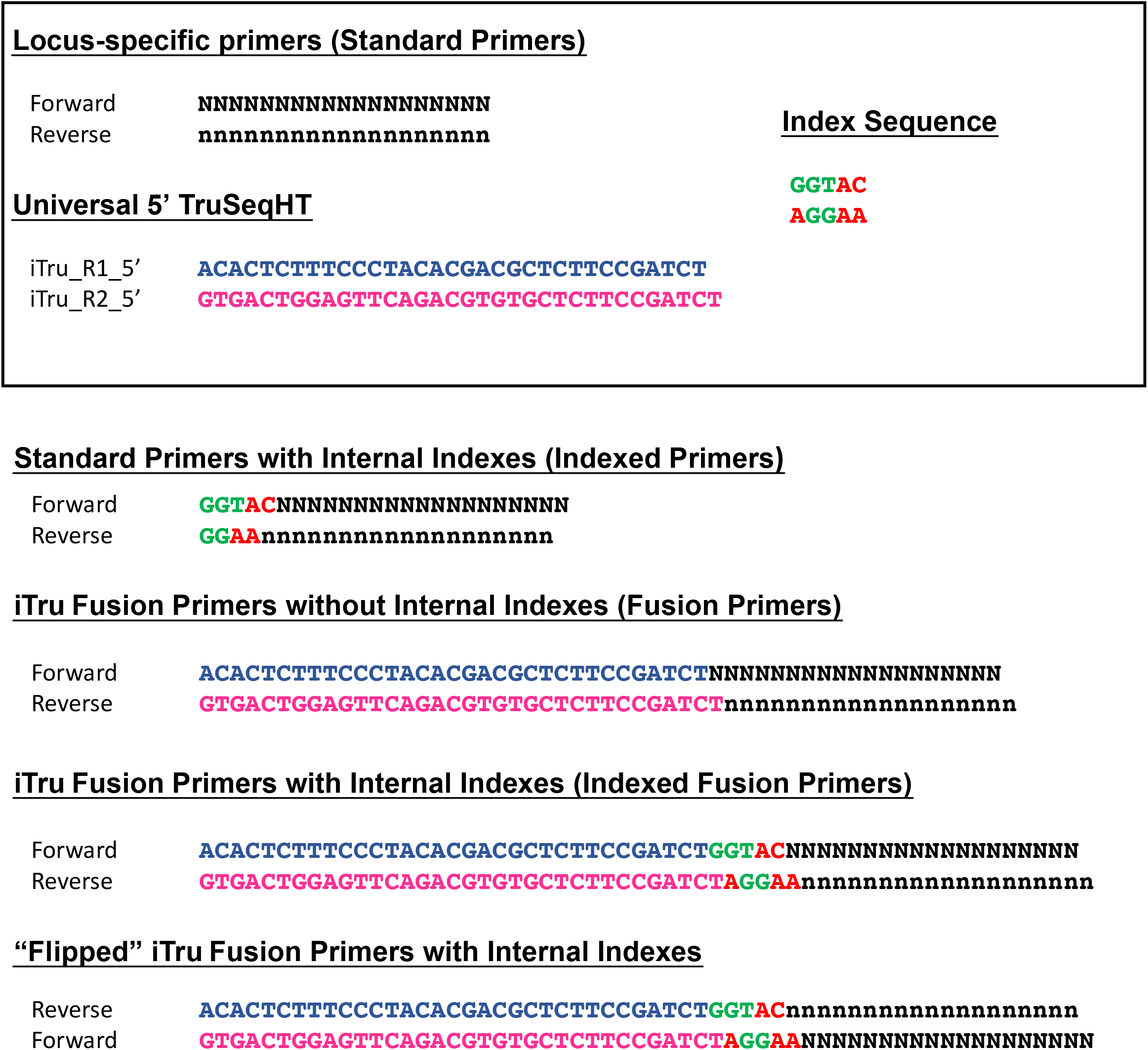
Examples of possible primer types (Table 3), including “flipped” fusion primers. Elements in the box are combined to form each of these various primer types, shown below the box. Standard locus-specific primer sequences are indicated by the letter “N”, in uppercase the forward primer and lowercase the reverse primer. Green and red nucleotide bases refer to unique index sequences. Blue and pink sequences are Read1 and Read 2 fusion sequences, respectively.

**Figure 3.**
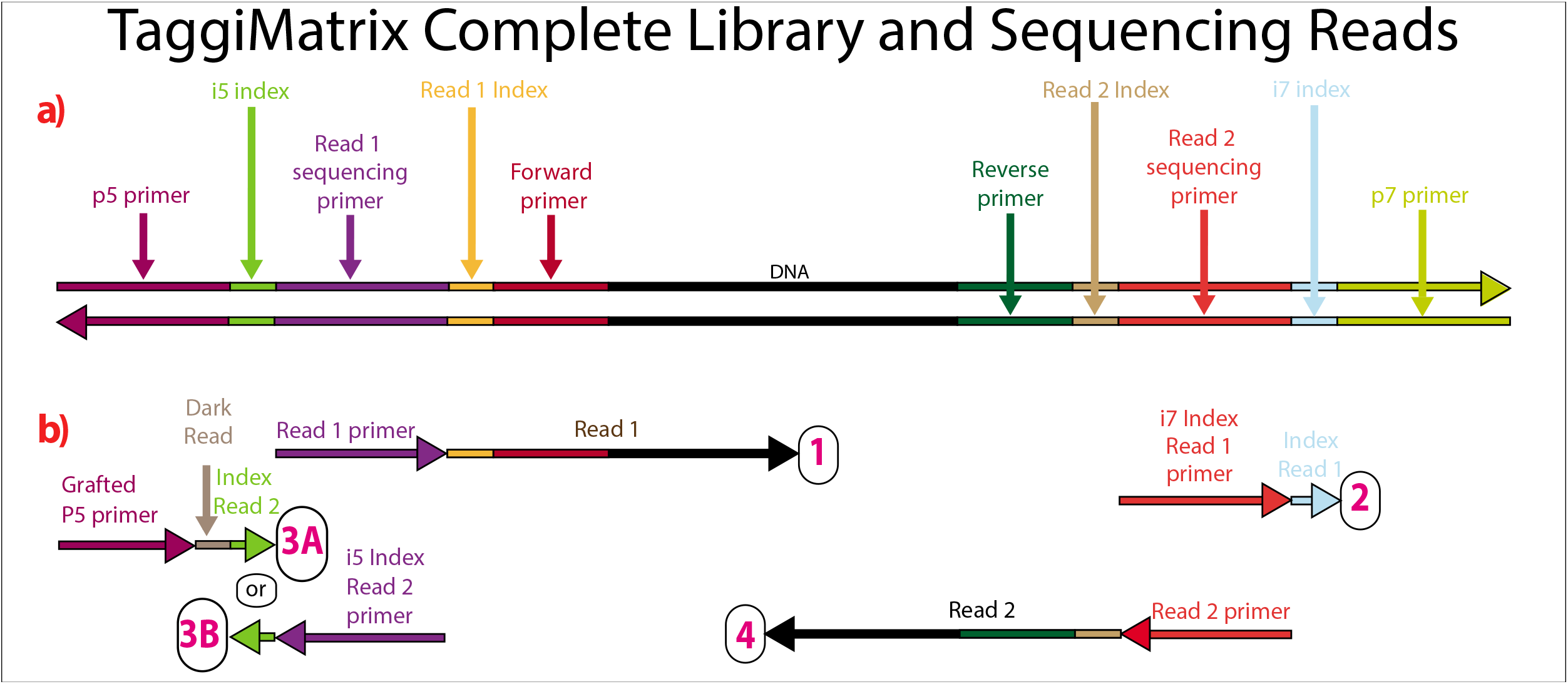
Sequencing reads that can be obtained from dual-indexed paired-end reads. a) Illustration of a double-stranded DNA molecule from a full-length amplicon library (i.e., following the limited-cycle round of PCR). Horizontal arrowheads indicate the 3’ ends. Labels on the double-stranded DNA indicate the function of each section, with shading to help indicate boundaries. b) Scheme of the four separate primers used for the four sequencing reactions that occur in paired-end dual-indexed sequencing and the reads that each primer produces (number in the circle). The four sequencing primers are added one at a time in the following order – Read1, Index Read1, Index Read2, and Read2. Vertical height indicates this order (top primer added first). 3A and 3B correspond to workflow A (NovaSeq™ 6000, MiSeq™, HiSeq 2500, and HiSeq 2000) and workflow B (iSeq™ 100, MiniSeq™, NextSeq™, HiSeq X, HiSeq 4000, and HiSeq 3000), respectively, of dual-indexed workflows on paired-end flow cells (Illumina 2018).

**Table 1.** Internal identifying index sequences. All indexes have an edit distance of ≥ 3. Upper case letters are the indexes; lower case letters add length variation to facilitate sequence diversity at each base position of amplicon pools (see text for details). For Illumina MiSeq and HiSeq models ≤2500, adenosine and cytosine are in the red detection channel, whereas guanine and thymine are in the green channel. Indexes and spacers have balanced red and green representation at each base position within each group of four indexes (i.e., count 1–4, 5–8, 9–12, 13–16, and 17–20).

| Index count | Index Label | Sequence | Length |
| --- | --- | --- | --- |
| 1 | A | GGTAC | 5 |
| 2 | B | cAACAC | 6 |
| 3 | C | atCGGTT | 7 |
| 4 | D | tcgGTCAA | 8 |
| 5 | E | AAGCG | 5 |
| 6 | F | gCCACA | 6 |
| 7 | G | ctGGATG | 7 |
| 8 | H | tgaTTGAC | 8 |
| 9 | 1 | AGGAA | 5 |
| 10 | 2 | gAGTGG | 6 |
| 11 | 3 | ccACGTC | 7 |
| 12 | 4 | ttcTCAGC | 8 |
| 13 | 5 | CTAGG | 5 |
| 14 | 6 | tGCTTA | 6 |
| 15 | 7 | gcGAAGT | 7 |
| 16 | 8 | aatCCTAT | 8 |
| 17 | 9 | ATCTG | 5 |
| 18 | 10 | gAGACT | 6 |
| 19 | 11 | cgATTCC | 7 |
| 20 | 12 | tctCAATC | 8 |

Although iTru and iNext primers facilitate quick and low-cost additions of dual-indexed adapters, this still requires a separate PCR reaction (but, see Discussion). Thus, when hundreds of amplicons are to be sequenced, it becomes economical to use additional internal indexes (Table 1) so that amplicons can be pooled prior to the use of iTru or iNext primers (Figs. 1 and 2). This approach should work with a wide variety of primers (e.g., Table 2). Such combinatorial indexing is designed to work in 96-well plate arrays but can be modified for other systems. Typically, eight indexed fusion forward primers (A–H) and 12 indexed fusion reverse primers (1–12) are designed and synthetized (File S1). Then, each DNA sample in each well of the 96-well plate can be amplified with a different forward and reverse primer combination (File S1, PCR_Set_up). These PCR products can be pooled and amplified using a similar combinatorial scheme with tagged universal iTru/iNext primers in the second PCR (Table 3), enabling the large-scale multiplexing of samples in one Illumina run (Table 4). Finally, because Illumina MiSeq platforms have documented issues in the quality of Read 2, particularly in GC-rich regions (Quail et al., 2012), fusion primers can be designed to swap forward and reverse primers with Read1 and Read2 fusions (e.g., R1Forward + R2Reverse, vs. R1Reverse + R2Forward; “flipped” primers) to account for this issue (Fig. 2). It is also possible to do replicate amplification with both sets of primers (regular and flipped), to significantly increase base diversity in amplicon libraries.

**Table 2.**
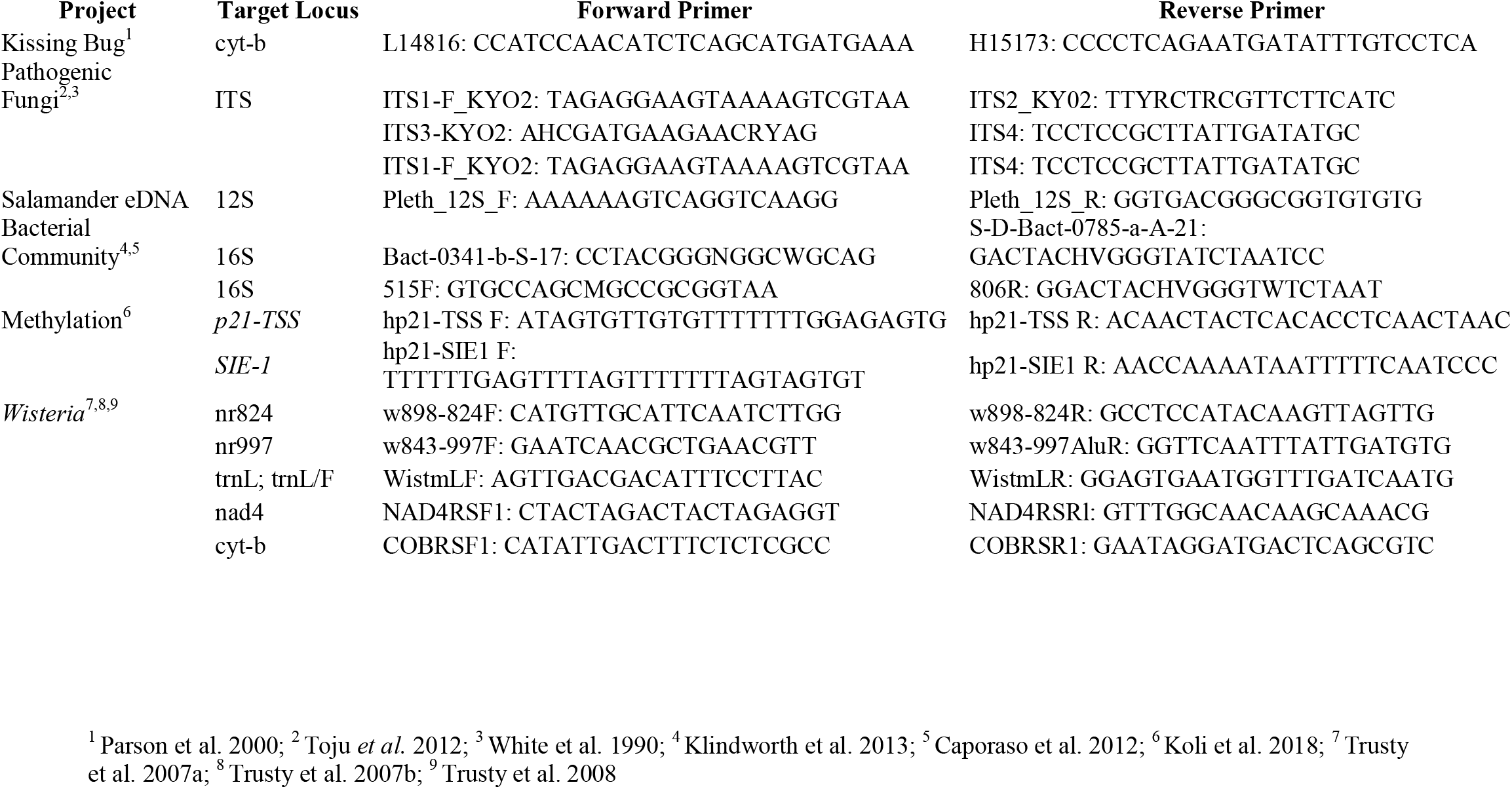
Primer pairs used in the example projects presented. Project, target locus, forward and reverse primer names and sequences, as well as the sources of the primer sequences are shown.

**Table 3.**
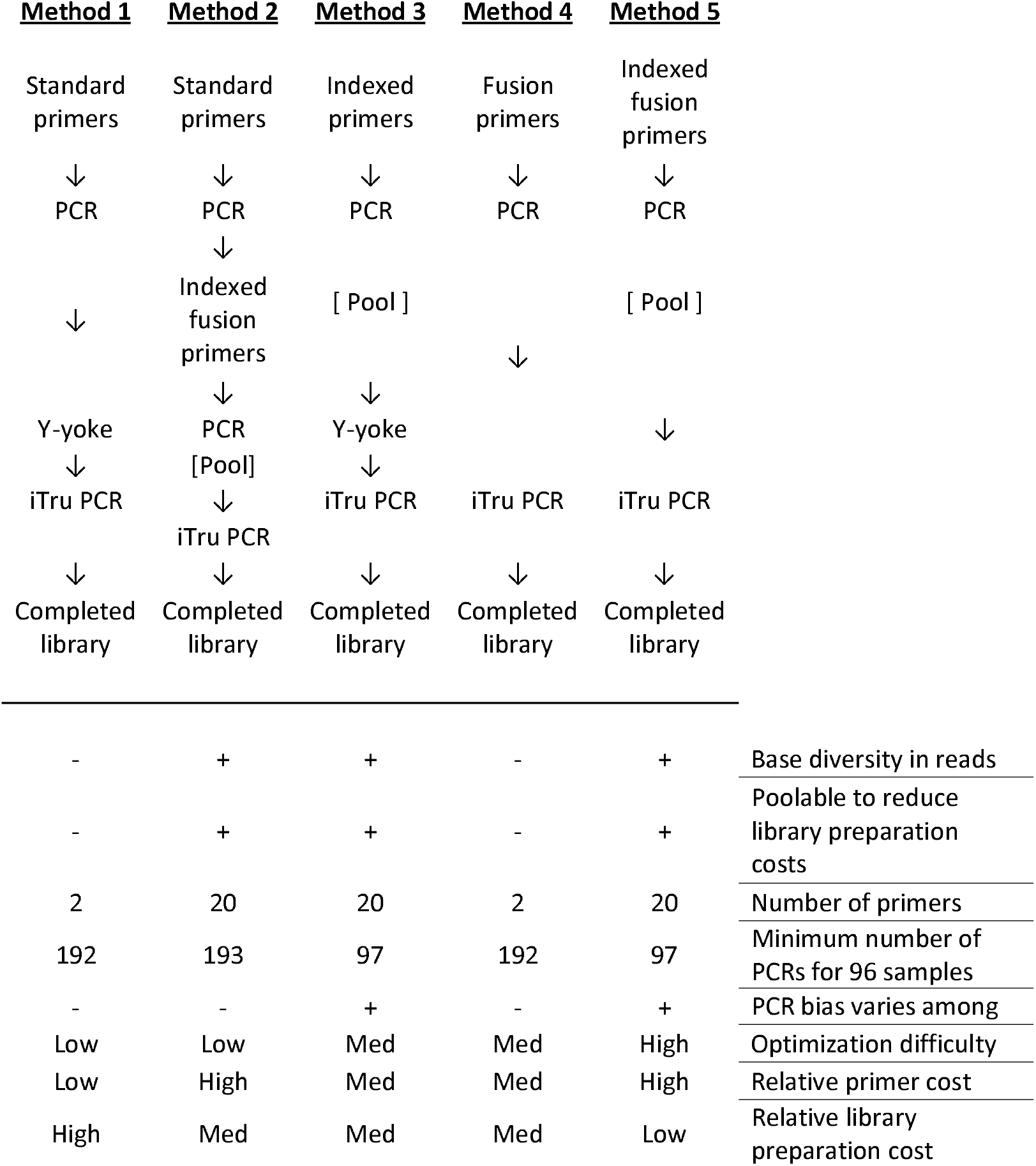
General strategies for producing and indexing amplicon libraries for Illumina sequencing. These examples use iTru primers, but as mentioned in the text, this can be implemented instead with iNext primers. Method 5 is illustrated below, but we are not including any dataset in the present manuscript that has implemented it (see Discussion). Note: this table does not include “flipped” primers.

**Table 4.**
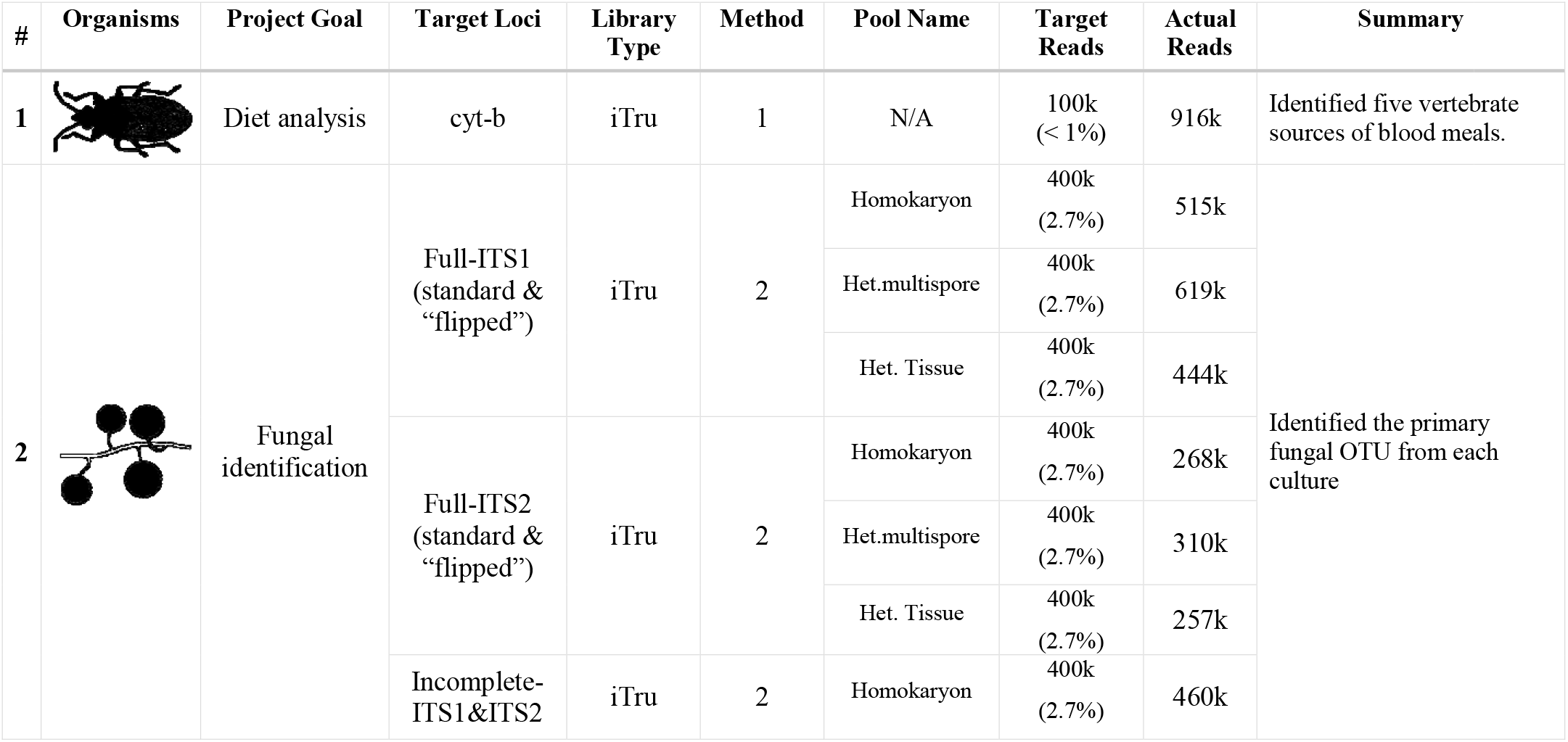
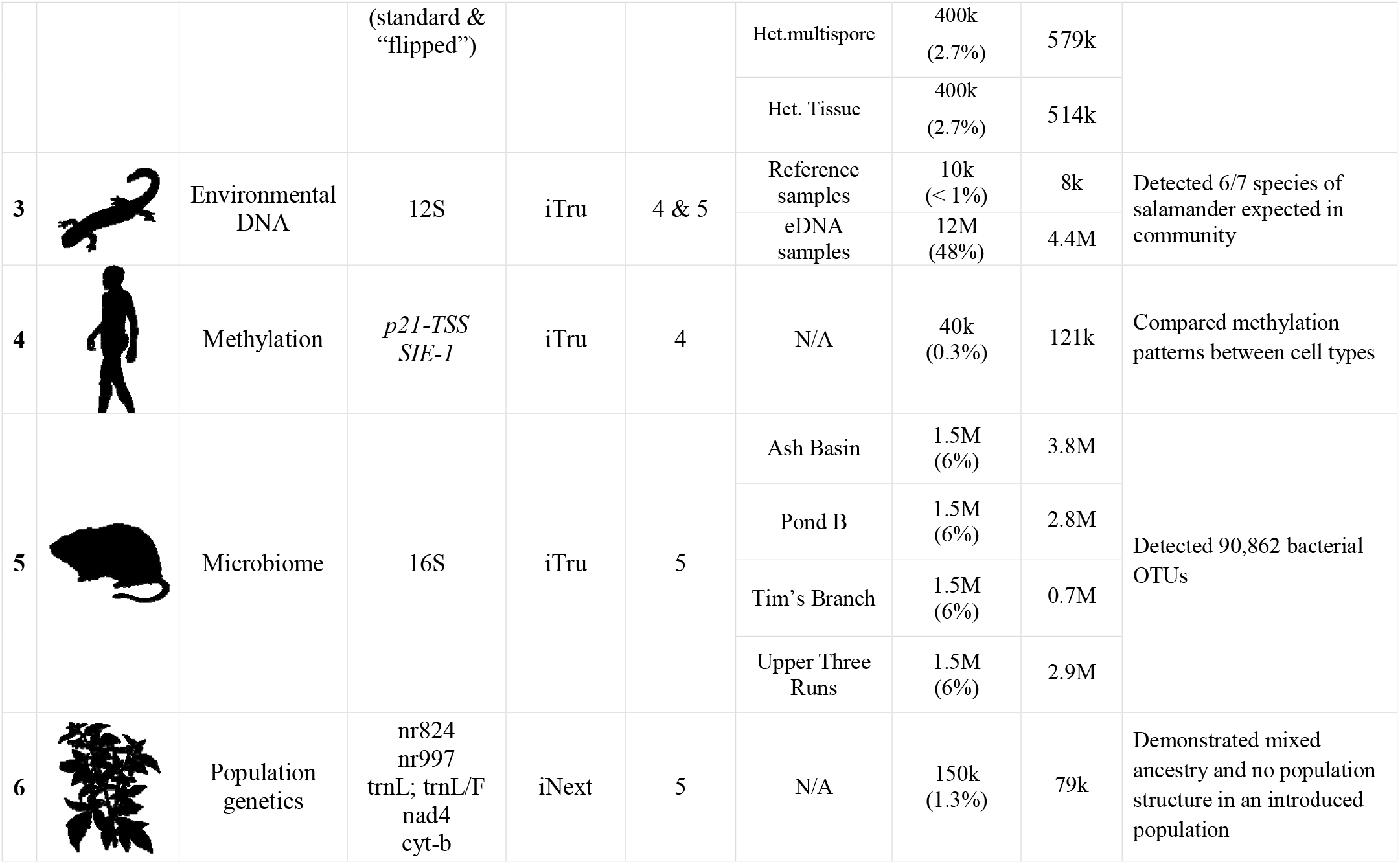
Detailed information for example projects presented to validate our approach. Summarized information for all example projects used to demonstrate Taggimatrix. The “Method” column refers to methods in Table 3; the “Target Reads” column cites the approximate number of reads per pool (i.e., not per individual sample) we targeted when pooling samples with other libraries. Note that these data were generated on many independent MiSeq runs. The kissing bug image is from Joseph Hughes (https://creativecommons.org/licenses/by-nc-sa/3.0/), and all other images are from PhyloPic 2.0 (Public Domain Dedication 1.0).

| # | Organisms | Project Goal | Target Loci | Library Type | Method | Pool Name | Target Reads | Actual Reads | Summary |
| --- | --- | --- | --- | --- | --- | --- | --- | --- | --- |
| 1 |  | Diet analysis | cyt-b | iTru | 1 | N/A | 100k (< 1%) | 916k | Identified five vertebrate sources of blood meals. |
| 2 |  | Fungal identification | Full-ITS1 (standard & “flipped”) | iTru | 2 | Homokaryon | 400k (2.7%) | 515k | Identified the primary fungal OTU from each culture |
|  |  |  |  |  |  | Het.multispore | 400k (2.7%) | 619k |  |
|  |  |  |  |  |  | Het. Tissue | 400k (2.7%) | 444k |  |
|  |  |  | Full-ITS2 (standard & “flipped”) | iTru | 2 | Homokaryon | 400k (2.7%) | 268k |  |
|  |  |  |  |  |  | Het.multispore | 400k (2.7%) | 310k |  |
|  |  |  |  |  |  | Het. Tissue | 400k (2.7%) | 257k |  |
|  |  |  | Incomplete-ITS1&ITS2 | iTru | 2 | Homokaryon | 400k (2.7%) | 460k |  |
|  |  |  | (standard & “flipped”) |  |  | Het.multispore | 400k<br>(2.7%) | 579k |  |
|  |  |  |  |  |  | Het. Tissue | 400k<br>(2.7%) | 514k |  |
| 3 |  | Environmental DNA | 12S | iTru | 4 & 5 | Reference samples | 10k<br>(< 1%) | 8k | Detected 6/7 species of salamander expected in community |
|  |  |  |  |  |  | eDNA samples | 12M<br>(48%) | 4.4M |  |
| 4 |  | Methylation | *p21-TSS*<br>*SIE-1* | iTru | 4 | N/A | 40k<br>(0.3%) | 121k | Compared methylation patterns between cell types |
| 5 |  | Microbiome | 16S | iTru | 5 | Ash Basin | 1.5M<br>(6%) | 3.8M | Detected 90,862 bacterial OTUs |
|  |  |  |  |  |  | Pond B | 1.5M<br>(6%) | 2.8M |  |
|  |  |  |  |  |  | Tim’s Branch | 1.5M<br>(6%) | 0.7M |  |
|  |  |  |  |  |  | Upper Three Runs | 1.5M<br>(6%) | 2.9M |  |
| 6 |  | Population genetics | nr824<br>nr997<br>trnL; trnL/F<br>nad4<br>cyt-b | iNext | 5 | N/A | 150k<br>(1.3%) | 79k | Demonstrated mixed ancestry and no population structure in an introduced population |

### TaggiMatrix applied case studies

We tested iTru primers designed as described above in five different experiments covering a wide range of experiments typically done in molecular ecology projects, and we tested iNext primers designed as described above in a single project (Table 4). In each experiment, we used at least two sets of primers: the first set (i.e., locus-specific fusion primers) generated primary amplicons, and the second set (i.e., iTru or iNext) converted primary amplicons into full-length libraries for sequencing (Fig. 3).

### iTru fusion primer experiments

For TruSeq-compatible libraries, we designed and synthetized locus-specific forward fusion primers, which started on the 5’ end with the Illumina TruSeq Read1 sequence (5’— ACACTCTTTCCCTACACGACGCTCTTCCGATCT—3’) for forward primers or the Illumina TruSeq Read2 sequence (5’—GTGACTGGAGTTCAGACGTGTGCTCTTCCGATCT—3’) for reverse primers; then included unique five nucleotide (nt) tags (Faircloth & Glenn, 2012) with variable length spacers (0–3 nt) to function as internal indexes (Table 1); and ended with locus-specific primer sequences (Fig. 2; Table 2). To assist with production of fusion primers and reduce errors, we have created and provided Excel spreadsheets (TaggiMatrix; File S1) and a web page (http://baddna.uga.edu/tools-taggi.html). With TaggiMatrix, users can simply input the names and sequences of the locus-specific primers, and all 22 (i.e., 2 non-indexed and 20 internally indexed) fusion primers and names are generated automatically. It is important to note that secondary structures or other PCR inhibiting characteristics are not checked by these tools (see Discussion). We then used the locus-specific fusion primers in a primary PCR, followed by a clean-up step and a subsequent PCR with iTru primers from *Adapterama I*. As an example, a general protocol for 16S amplification using TaggiMatrix can be found in File S2.

We used this approach for five projects (Table 4), each with slight modifications. First, we used primers targeting *cytochrome-b* to characterize the source of blood meals in kissing bugs; in this project, we first amplified DNA with standard primers, then ligated a y-yoke adapter to these products, and then amplified these products in an iTru PCR (Method 1 in Table 3). Second, we used primers targeting several portions of the ITS region, including “flipped” fusion primers, to identify fungal pathogens in tree tissues; in this project, we first amplified DNA with standard primers, then amplified these products with indexed fusion primers, and then amplified these products in an iTru PCR (Method 2 in Table 3). Third, we used primers targeting 12S to characterize plethodontid salamander communities from environmental DNA samples; in this project, we first amplified DNA with either internally indexed or non-indexed fusion primers and then amplified these products in an iTru PCR (Methods 4 or 5 in Table 3). Fourth, we used primers targeting two regions of the cyclin-dependent kinase inhibitor *p21* promoter to compare basal DNA methylation of *p21* promoter in two types of human cells; in this project, we first amplified DNA with non-indexed fusion primers and then amplified these products in an iTru PCR (Method 4 in Table 3; Kolli et al., 2019). Fifth, we used primers targeting 16S to characterize bacterial gut microbiomes in wild cotton mice (*Peromyscus leucopus*); in this project, we first amplified DNA with internally indexed fusion primers and then amplified these products in an iTru PCR (Method 5 in Table 3; File S2). Full methods describing the sample collection, DNA extraction, library construction (including detailed descriptions of pooling schemes), and data analysis are detailed in File S3.

### iNext fusion primer experiments

We generated libraries compatible with Nextera sequencing primers using the same approach as described above for TruSeq-compatible libraries, except that forward fusion primers started with Illumina Nextera Read1 sequence (5’— TCGTCGGCAGCGTCAGATGTGTATAAGAGACAG—3’), and reverse primers started with the Illumina Nextera Read2 sequence (5’— GTCTCGTGGGCTCGGAGATGTGTATAAGAGACAG—3’), and the second PCR used iNext primers from *Adapterama I* (Glenn et al., 2019). We have provided separate sheets within the TaggiMatrix Excel file (File S1) to facilitate the construction of iNext fusion primers.

We used this approach in one project. We used primers targeting one chloroplast locus, two mitochondrial loci, and two nuclear loci to perform a fine-scale population genetic analysis of the invasive vine *Wisteria*; in this project, we first amplified DNA with indexed fusion primers and then amplified these products in an iNext PCR (Method 5 in Table 3). Full methods describing the sample collection, DNA extraction, library construction (including detailed descriptions of pooling schemes), and data analysis are included in the File S3.

### Pooling, Sequencing, Analysis

The methods used for pooling, sequencing and analysis varied among the six projects (File S3), but some general approaches were consistently employed. Amplicon library pools from each of the six projects were pooled with additional samples and sequenced at different times on Illumina MiSeq instruments. The sizes of the amplicons were determined from known sequence targets and verified by agarose gel electrophoresis and known size-standards. We quantified purified amplicon pools using Qubit (Thermo Fisher Scientific Inc, Waltham, MA). We then input the size, concentration, and number of desired reads for amplicon sub-pools and all other samples or sub-pools that would be combined together for a sequencing run into an Excel spreadsheet to calculate the amount of each sub-pool that should be used (an example file of our pooling guide can be found in File S4). We targeted total proportions ranging from <1% to 44% of the MiSeq runs (Table 4). We used v.3 600 cycle kits to obtain the longest reads possible for four of the projects and v.2 500 cycle kits for two of the projects, which reduces buy-in costs when shorter reads are sufficient.

Following sequencing, results were returned via BaseSpace or from demultiplexing the outer indexes contained in the bcl files using Illumina software (bcl2fastq). Following demultiplexing of the outer indexes, we used Mr. Demuxy (https://github.com/lefeverde/Mr_Demuxy; File S5) or Geneious^®^ to demultiplex samples based on internal indexes.

Downstream analyses varied according to the goals of each project and further details are found in File S6. In brief, after demultiplexing, we cleaned raw sequencing data from each project by trimming primers and quality-filtering. Then, we compared sequences from project 1–4 against relevant databases to identify OTUs. For projects 5–6, we mapped reads to appropriate reference sequences. For project 5, we extracted methylation profiles, whereas for project 6, we identified sequencing polymorphisms among genes and individuals. Additional details about each project are presented in Supplemental File S3.

## Results

We used five methods that take advantage of iTru or iNext indexing primers developed in *Adapterama I* in six exemplar amplicon sequencing projects. These projects illustrate the range of methodological approaches that can be used to overcome challenges of amplicon library preparation and fulfill most of the characteristics of an ideal amplicon library preparation system.

In all but one project (Table 4, project 1), we designed fusion primers to generate amplicons that can be amplified by iTru5 and iTru7 (or iNext5 and iNext7) primers to create full-length Illumina TruSeq (or Nextera) libraries. The indexed fusion primers utilize 20 (i.e., 8 + 12) internal identifying sequences with an edit distance ≥ 3 (Table 1) to create up to 96 internally dual-indexed amplicon libraries which were used individually or pooled for additional outer indexing by iTru5 and iTru7 (or iNext5 and iNext7) primers. Sequential PCRs that start with internally indexed primers create quadruple-indexed amplicon libraries that achieve our design goals of cost reduction, facilitation of large-scale multiplexing, increased base-diversity for Illumina sequencing, and maximization of efficiency of library preparation.

In our project characterizing the blood meals of kissing bugs (Table 4, project 1), we obtained an average of 116,902 reads for each sample and identified a total of five unique vertebrate species as the source of the blood meals. In our project identifying fungal pathogens in tree tissues (Table 4, project 2), we obtained an average of 436,825 reads per pool (i.e., 96 samples) and characterized the diverse fungal communities found in these samples. In our project characterizing plethodontid salamander communities from environmental DNA samples (Table 4, project 3), we obtained an average of 163,555 reads for each PCR replicate and identified reads matching 6/7 species expected to be present in the streams. In our project comparing basal DNA methylation of *p21* (Table 4, project 4), we obtained approximately 10,000 reads per sample and detected differences in methylation of CpG sites between embryonic kidney cells and human proximal tubule cell (Kolli et al., 2019). In our project characterizing bacterial gut microbiomes (Table 4, project 5), we rarified to 15,000 quality-filtered reads per sample and identified an average of 3,847 OTUs per sample. In our project focused on the fine-scale population genetic analysis of *Wisteria* (Table 4, project 6), we obtained an average of 1,697 reads per sample and discovered little evidence of population structure among samples. Variation in the average number of reads among projects reflects the intentional allocation of reads when pooling with genomic libraries for sequencing; for example, we pooled plates of libraries for the fungal pathogen project in relative quantities intended to generate approximately 4,000 reads per sample. Variation in the number of reads among samples within a given project likely reflects quantification error and variation in input DNA quantity and quality. Full results and associated figures for each project are detailed in File S3.

The costs associated with each method vary significantly, and which approach has the lowest cost depends on the number of samples processed (Fig. 4: note axis scales are not linear; Table 5; File S6). Methods 1 and 4 have the lowest buy-in cost, but the cost of library preparations are fixed, rather than decreasing as the number of samples increases. The constant cost per sample is due to the need for individual second round PCRs (e.g., iTru5/7). The other methods allow pooling of samples prior to second round PCR, which reduces costs. Because Method 1, with no use of fusion primers (non-indexed/indexed), has the highest library preparation costs per sample, it quickly becomes the most expensive method, more than doubling the cost of most other methods with as few as 96 samples. Method 4 remains economically reasonable for processing one or two plates of samples but becomes less reasonable as more plates of samples are used. Method 2 is never economically best, but it is sometimes necessary to achieve sufficient amplification to construct the desired libraries. Thus, Method 2 is only viable when the other methods fail. Method 3 has a moderate buy-in cost and the second-lowest cost per sample for large numbers of samples. Also, Method 3 has the lowest cost when ≤11 plates of samples will be processed, though the cost is very similar to Method 5 after ≥2 plates of samples are processed. Method 5 has the second highest buy-in costs, but the lowest costs per sample when large numbers of samples are processed. Method 5 is optimal when >12 plates of samples are processed. Because Methods 3 and 5 are similar in cost after a few plates of samples are processed, other considerations, such as workflow and personnel costs, are likely to drive decisions about the optimal method rather than the costs of reagents.

**Figure 4.**
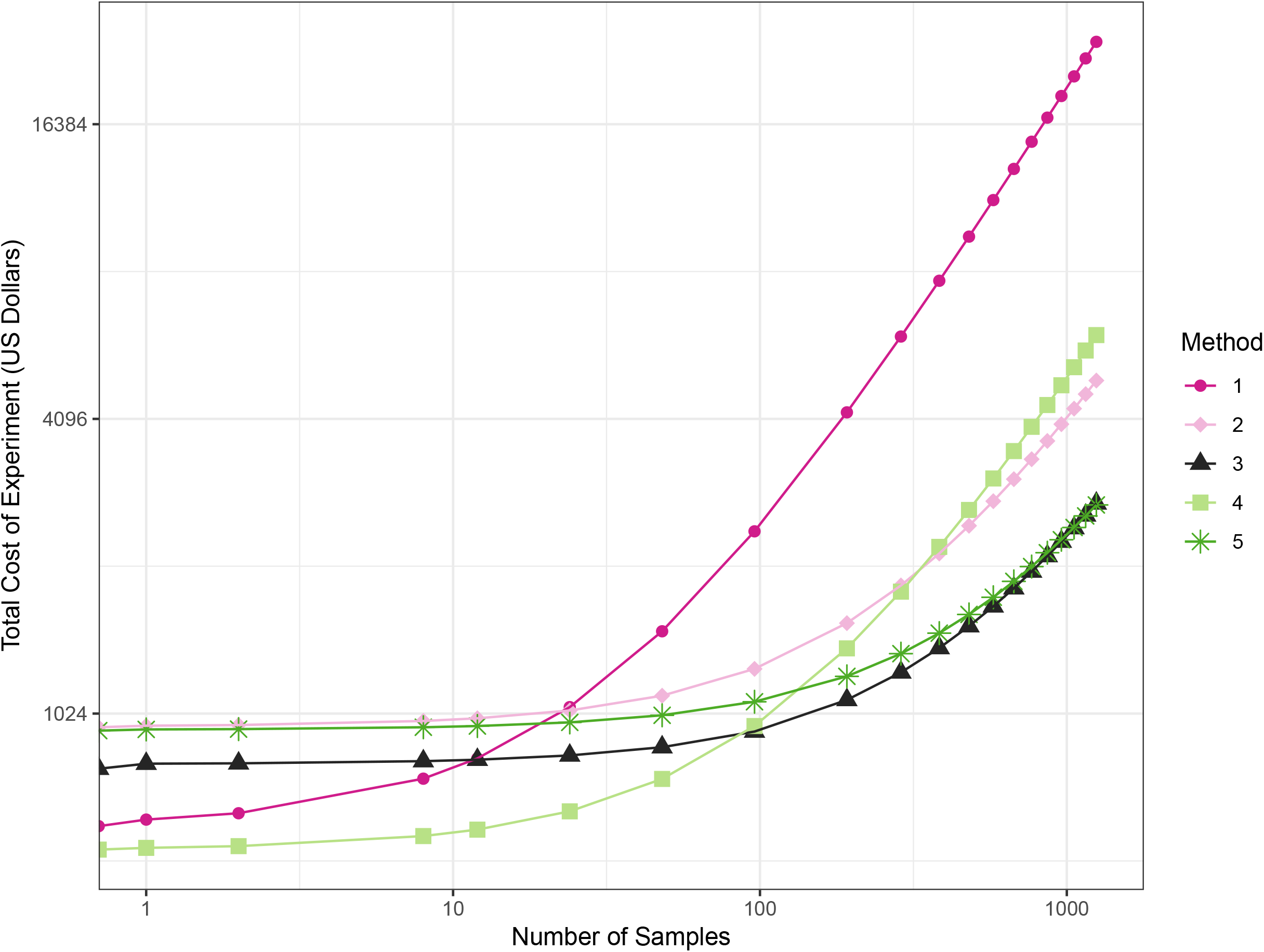
Total cost of experiments across the five methods given a number of samples. Line plot of price of each method according to the number of samples. The starting point in the X-axis (x=0) represents the buy-in cost of oligos.

**Table 5.**
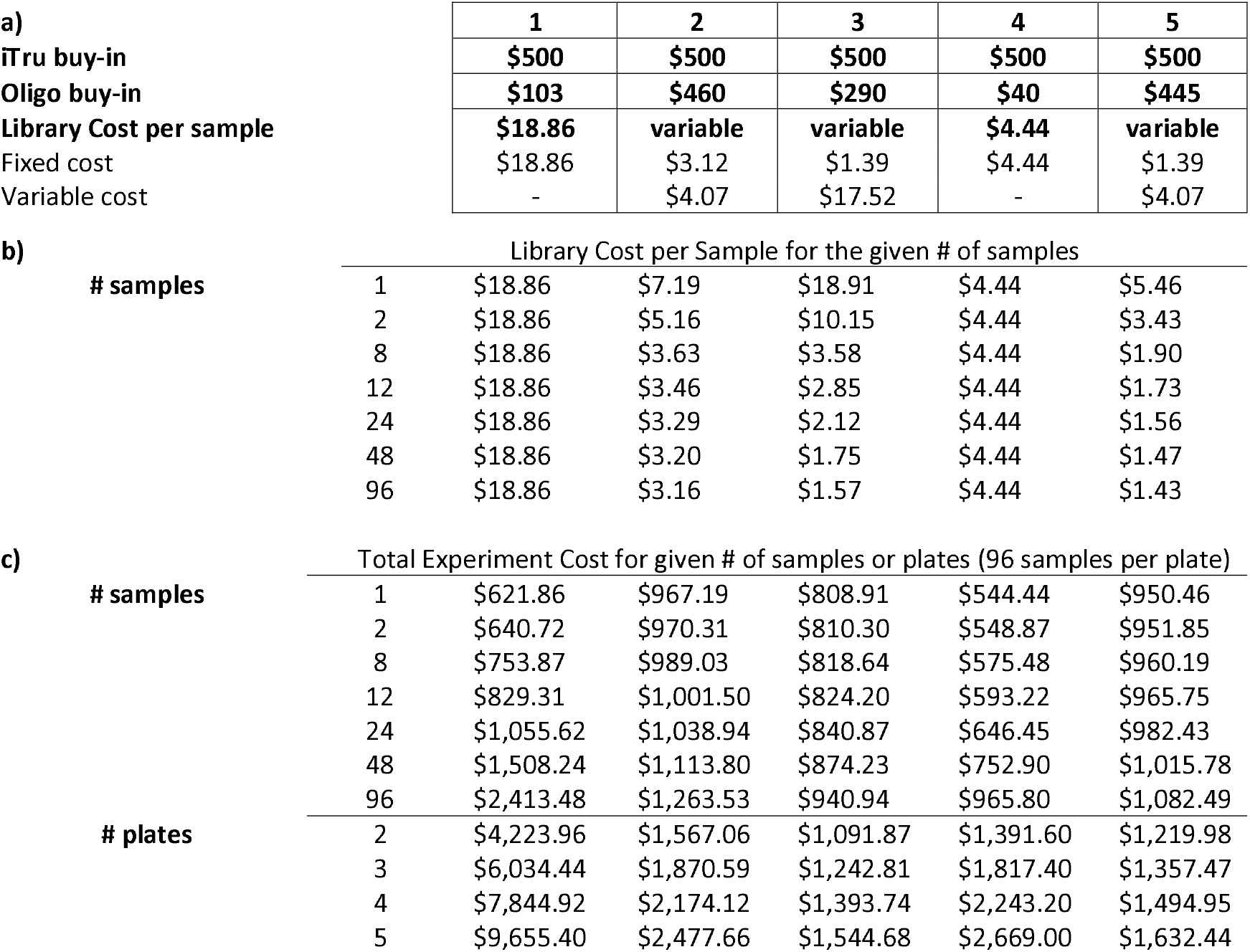
Oligos and iTru buy-in, and library prep costs among methods. Costs associated to the implementation of the different methods. In segment **a)** we present buy-in cost of oligos and iTru primers and cost per sample of library prep which consists of both, fixed and variable costs depending on pooling at early stages. Segment **b)** is the cost of library prep (no considering primers/adapters) per sample given a number of samples. Segment **c)** is the total experimental cost of primers/adapters and library prep according to the number of samples in the experiment, the first section is in term of number of samples, the second section is in terms of plates, each plate consisting of 96 samples. Cost for iTru are calculated list prices of aliquots from baddna.uga.edu. Costs for ‘oligos’ are calculated using list prices from Integrated DNA Technologies (IDT; Coralville, IA). Other costs are from listed prices from various vendors by Jan 2019. Please view File S1 and S6 for additional details on price calculations and also to review total prices of experiment given a number of samples.

## Discussion

In *Adapterama I*, we introduced a general approach to reduce the cost of genomic library preparations for Illumina instruments. Here, we made extensive use of the iNext and iTru primers described in *Adapterama I* and show that these can also be used to facilitate amplicon library construction at reduced cost with increased flexibility. As we did in *Adapterama I*, we focused mostly on iTru to simplify our presentation of the method, but iNext works identically in most situations.

Although we focused on Illumina, many of these approaches can be extended to other platforms following the design principles described here (e.g., use primers from sheet ITS_10nt_5’tags in File S1 following Method 3). For platforms that sequence individual molecules (e.g., PacBio and Oxford Nanopore), there is no advantage to variable-length indexes and negligible penalty for longer indexes, but there are significant informatic advantages to equal-length indexes. Thus, for many other platforms, it will be better to use longer indexes of equal length.

In general, TaggiMatrix Method 5 achieves our design goals, in that it: 1) uses the universal Illumina sequencing primers; 2) minimizes costs (as little as $2.20 per library, i.e. Method 3 when prepping 1,248 samples in thirteen pools, Figure 4, File S6); 3) minimizes time and equipment needed for library preparations; 4) minimizes buy-in costs through the use of a limited number of fusion primers and universal iTru7 and iTru5 primers; 5) eliminates error-prone ligation steps; 6) allows for > thousands of samples to be pooled and run simultaneously; 7) allows users to vary amplicon representation from tiny to large fractions of a sequencing run (up to 91% has been validated for other projects, data not shown); 8) supports creating millions of samples (8 × 12 × 384 × 384 = 14,155,776) that can be tracked and multiplexed through quadruple-indexing. TaggiMatrix Method 3 shares nearly all of these advantages; per sample costs are a few cents more and ligation of a universal stub onto the amplicon pool is maintained.

Similar to other *Adapterama* applications, TaggiMatrix offers several methods for combinatorial and hierarchical indexing of samples (Table 3), allowing users to optimize various criteria. For example, different indexes can be used at any combination of the four index positions in the TaggiMatrix library (Fig. 3). By using inner indexes in combination, 20 (8 + 12) indexes can be used to identify 96 (8 x12) samples. By using inner and outer indexes hierarchically, 40 (8 + 12 + 8 + 12) indexes can identify 9216 (8 × 12 × 8 × 12) samples. By using two sets of iTru5 and iTru7 primers, 36,864 (8 × 12 × [8 + 8]x[12 + 12]) samples can be identified. Varying indexes at all index positions is the most economical way to tag samples, especially as the number of samples increases (Table 6). By combining a single set of 20 (8 + 12) fusion primers with the full set of 384 iTru5 and 384 iTru7 primers from *Adapterama I* (Glenn et al., 2019), a total of 14,155,776 (8 × 12 × 384 × 384) samples can be multiplexed.

Our methods address the issue of base diversity through the incorporation of indexes with variable-length spacers that allow for diversity at each base position. This strategy is based on independently originating ideas implemented at the Broad Institute, our lab and others, such as the system developed by Fadrosh et al. (2014) where they introduced “heterogeneity spacers” for sequencing amplicons out of phase. Longer spacers (e.g., 0–7 nt) are advantageous over shorter spacers to compensate for longer repeats in the target amplicons. Mononucleotide repeats are particularly problematic in terms of base diversity. Mononucleotide repeats of ≥5 bp will not be addressed by our short spacers (Table 1). Because Illumina reads are of set length, longer spacers decrease the total amount of useful sequence obtained for downstream analyses. Thus, there is a trade-off in how long the heterogeneity spacers should be. Here, we implement a 0–3 nt long heterogeneity spacers, although this could be easily tuned to 0–7 nt for forward primers and 0–11 nt for reverse primers, to accommodate any researcher’s preferences and mononucleotide repeats known to occur in the target sequences.

Our approach does not deal with the limitation of read-length on Illumina platforms. For long amplicons where complete sequencing is desired, it is possible to construct shotgun libraries from the longer amplicons (e.g., using Illumina Nextera XT, Kapa Biosystems Hyper Prep Plus, NEB Ultra II FS or many other commercial kits). The methods used in *Adapterama I* may be helpful in those cases. Such libraries can take advantage of the reduced costs per read on higher capacity instruments. It is also possible to design internal locus-specific fusion primers that recover the entire desired DNA region through independent PCRs. It is important to note, however, that the recent introduction of the PacBio Sequel II along with sequencing chemistry v.6 makes circular consensus sequencing of long amplicons on PacBio an economically reasonable approach. Thus, use of the longer consistent-length indexes noted above to create amplicon pools for PacBio is likely to be increasingly attractive as their platform continues to improve.

TaggiMatrix provides an easy way to create indexed fusion primers for convenient ordering at any oligo vendor of your choice. However, the current web page and spreadsheets do not perform quality control of the primer sequences generated. Thus, before ordering, it is important to validate the fusion primers to ensure hairpins, dimers and other secondary structures that inhibit PCR are not created. Several programs exist to validate the primers designed and these should be used before ordering. It is also generally recommended that a small number of fusion primers should be obtained and tested prior to investing large batches of long fusion primers. When deciding on the best method to use (i.e., Methods 1–5), the number of samples, reagent cost, and time available to optimize the primers should be considered (Fig. 5).

**Figure 5.**
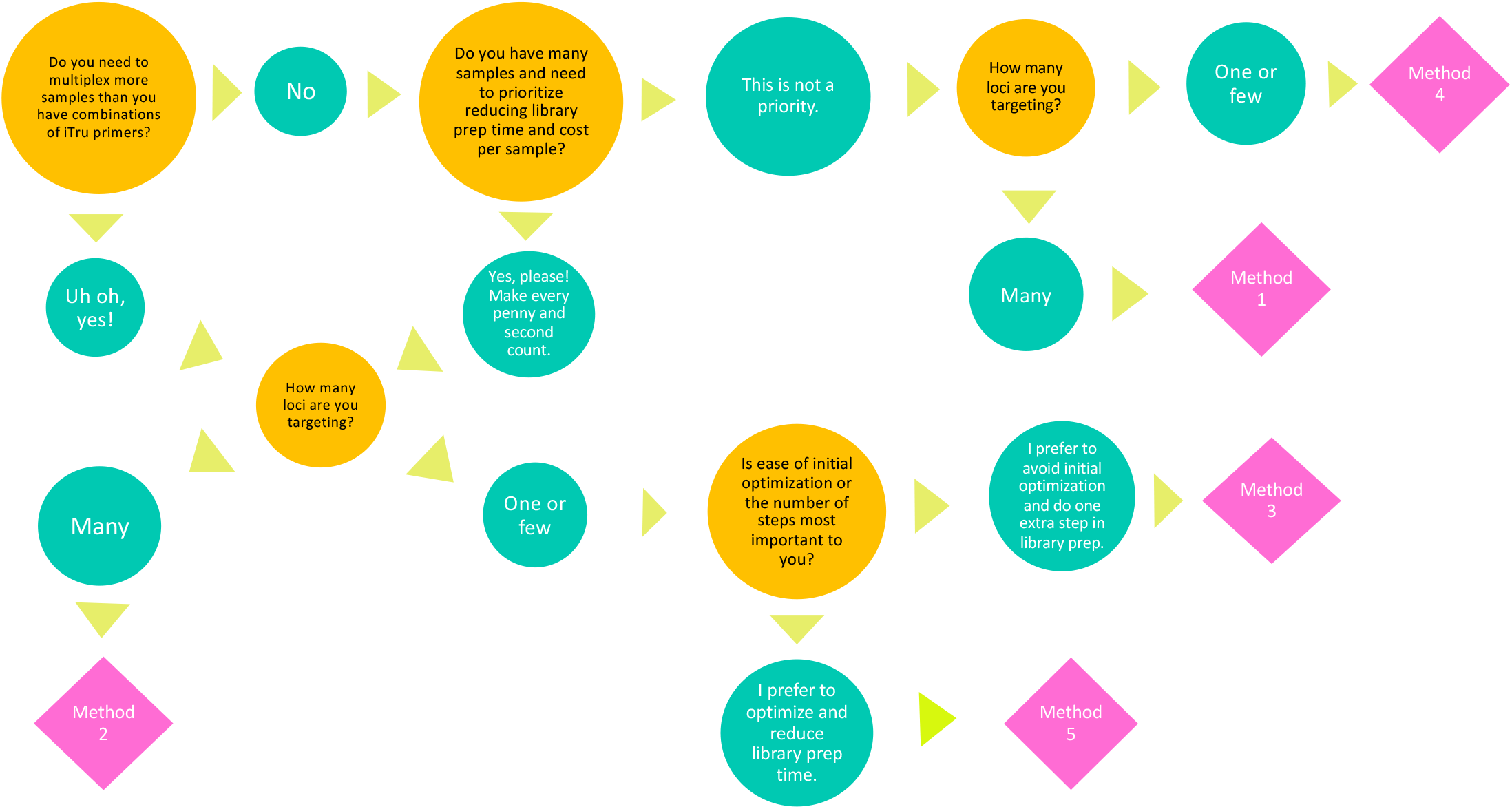
Decision tree to select the best fitting method according to the experiment goals and budget. Guide of choices to drive an informed decision over the method for amplicon sequencing that may be fit the best for your lab/research/experiment goals.

While developing adapters and primers to make multiple libraries that will be pooled and sequenced, it is important to determine if the primers with different indexes have biased amplification characteristics. This can be accomplished by testing all primers via quantitative PCR using a common template pool to ensure that each primer was synthesized, aliquoted, and reconstituted successfully and has similar amplification efficiency. In practice, however, it will not be economical or necessary to conduct such rigorous quality control for many projects. It is important to note that because sequencing reads are so cheap (~10,000 reads per $1 USD for PE300 reads on a MiSeq), being off by thousands of reads per sample is less expensive than precise quantification, especially when personnel time for such quantification is considered. Thus, it will often be less expensive to subsample reads from overrepresented samples and/or simply redo the small proportion of samples that do not generate a sufficient number of reads. Another common concern with amplicon library preparation methods involving PCR is the introduction of bias due to PCR duplicates. Our method can be modified to incorporate 8N indices similar to how we addressed this issue with RADcap libraries (Hoffberg et al., 2016). It is also possible to use internal N indices of any length desired as molecular identifiers (i.e., Jabara et al., 2011; Kou et al., 2016). These modifications, in conjunction with long-amplicon sequence on other platforms is worthy of further work.

## Conclusions

In summary, we demonstrate how several variants of TaggiMatrix solve common challenges for amplicon sequencing on NGS platforms. Our methods can be implemented in projects from a wide array of disciplines such as microbial ecology, molecular systematics, conservation biology, population genetics, and epigenetics, and we encourage others to further develop the tools we provide for solving additional challenges posed by these applications.

## Acknowledgements

We thank our colleagues at the Georgia Genomics and Bioinformatics Core. We thank our many colleagues at UGA and elsewhere who tested early versions of these protocols over the past five years. We thank John Maerz for his help collecting environmental DNA samples. We thank Bradley Brown for his technical help on Bismark.

---

Supplementary Figure S1

**Diagram of full-length amplicon TaggiMatrix library product**

Double stranded amplicon library product after implementation of TaggiMatrix. Indication tags and indexes incorporated through the use of Fusion primers and iTru/iNext primers, respectively.

Supplementary Figure S2

**Detailed illustration of the components on one of the possible designs (Method 5) to construct TaggiMatrix amplicon libraries**

First, locus specific fusion primers with tags are used to amplify the target DNA region. From this step pooling is possible thanks to the presence of indexes. Then library amplification with the use of iTru univer primers with indexes that allows pool labeling and incorporation of Illumina platform oligos (P5 and P7).

Supplementary File S1

**TaggiMatrix spreadsheet**

Excel spreadsheet demonstrating the step-by-step process to create indexed fusion primers with TaggiMatrix. The first sheet (Introduction) is an introductory explanation of how the document works. The second, third, and fourth sheets (…iTru_Fusions) are examples of the creation of indexed fusion primers for 16S, cyt-b and COI universal primers, respectively. The fifth sheet (iNext_&_iTru_Primers) is a list of the universal primer sequences and prices. The sixth and seventh sheets (…Order_Sheet) are examples of how to fill the order form to fill plates with primer sets. The eighth sheet (PCR_Setup) indicates how to combinatorically layout the primers for a 96-well plate. The ninth, tenth, and eleventh sheets (…Tags…) list the index sequences that are incorporated to the fusion primers, their spacers, and examples.

Supplementary File S2

**TaggiMatrix protocol for 16S amplicon library prep**

Step-by-step library construction for 16S libraries with indexed fusion primers.

Supplementary File S3

**Supplementary methods and results for TaggiMatrix example datasets**

A detailed guide through the methods, results, and discussion of sequence analyses from TaggiMatrix data generated for each example dataset presented in this manuscript.

Supplementary File S4

**TaggiMatrix video: what is happening inside the tube?**

This presentation demonstrates the key features of TaggiMatrix, including how the combinatorial indexing is performed in a plate.

Supplementary File S5

**Demultiplexing Internal Indexes Using Mr. Demuxy**

Guide of how to run Mr. Demuxy to demultiplex using internal indexes amplicon data in fastq format.

Supplementary File S6

**Price calculator among methods presented for amplicon sequencing**

Excel spreadsheet with calculations of oligos and reagents costs for library prep among the five methods presented in *Adapterama II*. Users can modify values according to their particular vendors, number of samples, and number of pools, to have an estimate of the price per sample and the price of the experiment.

The authors declare competing interests. The EHS DNA lab provide oligonucleotide aliquots and library preparation services at cost, including some oligonucleotides and services that make use of the adapters and primers presented in this manuscript (baddna.uga.edu). The information we present allows all researchers to synthesize the oligonucleotides at any vendor of their choice, follow or modify the library preparation techniques we have included, and freely publish results simply with proper attribution of this paper and Illumina^®^™. Any use of trade, firm, or product names is for descriptive purposes only and does not imply endorsement by the U.S. Government.

This work was partially supported by: DEB-1242241, DEB-1242260, Dimensions of Biodiversity DEB-1136626, DEB-1146440, Graduate Research Fellowships DGE-0903734 and DGE-1452154, and Partnerships for International Research and Education (PIRE) OISE 0730218 from the U.S. National Science Foundation.

