## Supplementary figures and images for "Adapterama II: Universal amplicon sequencing on Illumina platforms (TaggiMatrix)"

### Fig. S2

#### TaggiMatrix Library Components

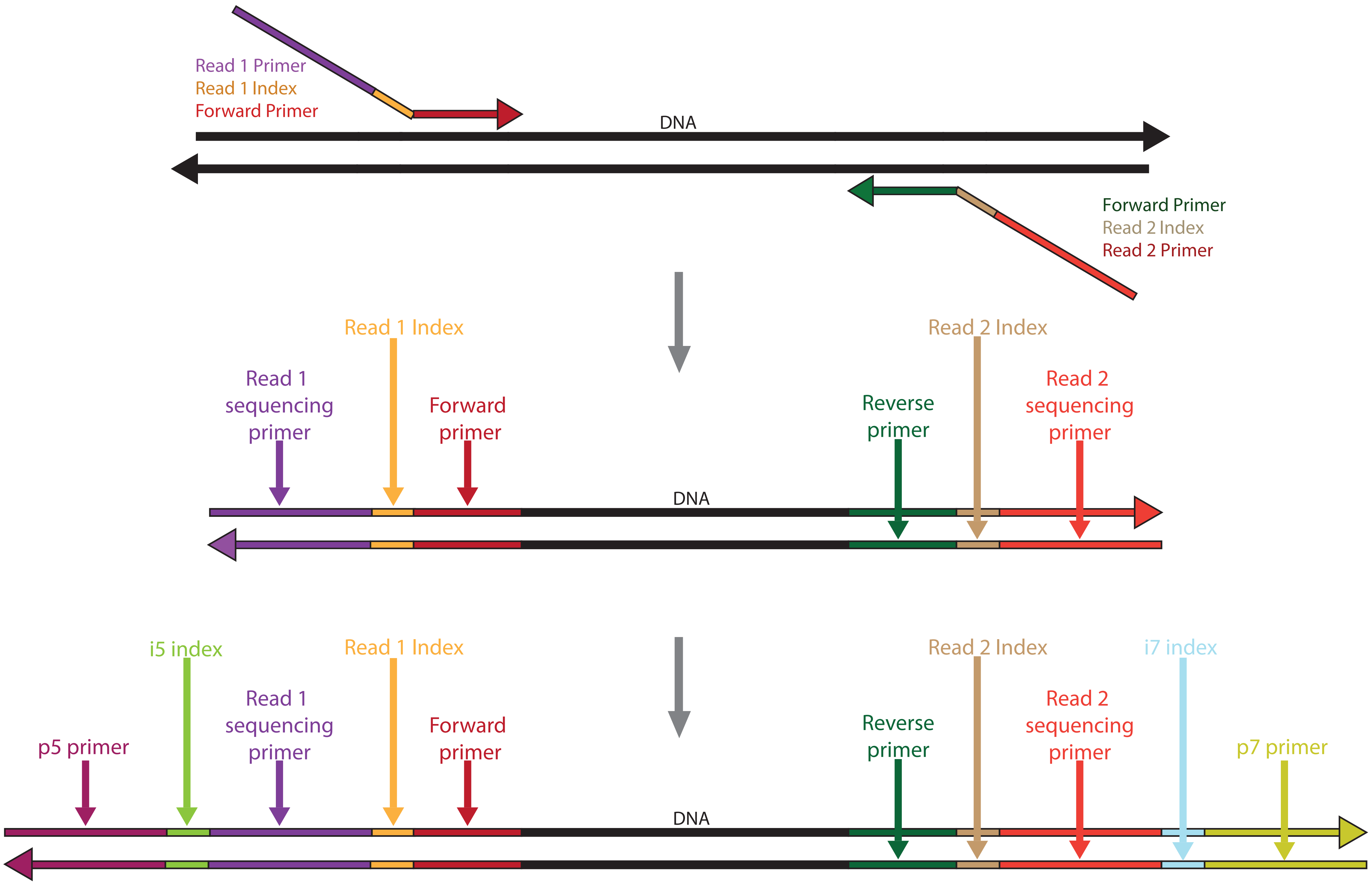

### Figs. S1

#### Index Positions in TaggiMatrix Complete Library

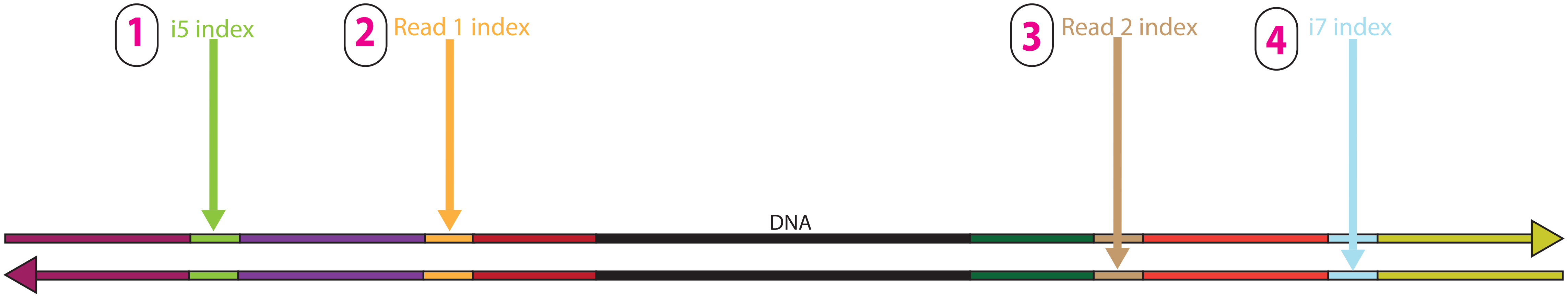
