## Supplementary material for "Adapterama II: Universal amplicon sequencing on Illumina platforms (TaggiMatrix)": File S2

**TaggiMatrix 16S PCR Protocol using fusion-indexed primers**

1. PCR recipe when using fusion-indexed 16S primers:

Create the master mix with following recipe per sample (alter amount of water or DNA based on sample), add extra 10% of volume of each reagent:

10.75 μL H2O

5.0 μL 5x Kapa HiFi Buffer (Kapa Biosystems, Wilmington, MA)

0.75 μL dNTP’s (10 mM stock from Kapa kit)
0.5 μL Kapa Hifi DNA Polymerase (1 unit/μL from Kapa kit)

17 μL Master Mix

2. Vortex master mix thoroughly and centrifuge mix.
3. Pipette 17 μL of master mix into strip-tubes or 96-well plate
4. Add 5 μL of sample DNA (concentration normalized) to each strip-tube or plate well to reach a 22 μL volume
5. Arrange 16S fusion-indexed primers in order of 1-12 and A-H (as shown below)
6. Using multichannel pipet place 1.5 μL of Forward (1-12) fusion-indexed primers 5 μM into each row of samples horizontally
7. Using multichannel pipet place 1.5 μL of Reverse(A-H) fusion-indexed primers 5 μM into each column of samples vertically to obtain a final volume of 25 µL.

Here a graphical representation:

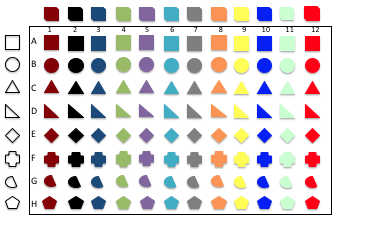

8. Vortex and centrifuge.

Place plate/samples into thermocycler using the following settings, 98 °C for 3 min, followed by 20-25 cycles at 98 °C for 30 sec, 55 °C for 30 sec, and 72 °C for 30 sec; and final extension at 72 °C for 5 min, holding after at 4 °C. [Note: Adjust the number of cycles based on the amount of input DNA, more cycles if less input DNA was used.]

9. Run PCR product in a 1.5% agarose gel. Bands should appear around 500bp region of ladder.

Note: If any sample did not amplify, place them back in the thermocycler for few additional cycles.

10. Pool 5 µL of each sample PCR product by pipetting with a 12-chanels multichannel vertically into a 12-tubes-strip-tube. Save the rest of product properly labeled as a backup. The total volume in each tube should be (5 µL X 8 A-H) = 40 µL. Then transfer each 40 µL to a 1.5 mL microtube to obtain a final pool of (40 µL X 12 (1-12)) = 480 µL
11. Add SpeedBeads (Thermo-Scientific, Waltham, MA, USA) to 1:1 ratio, purify as normal and resuspend in 100 µL of TLE.
12. PCR recipe to add complete Illumina adapters & indexes (three per plate):

10.0 μL 5x Kapa HiFi Buffer

1.5 μL dNTP’s (10 mM stock from Kapa kit)

17.5 μL dH_2_O (to make final total volume 50 µL)

1.0 μL Kapa HiFi DNA Polymerase (1 unit/μL from Kapa kit)
5.0 µL iTru5 Primer(s) (5µM -> 0.5 µM final)

5.0 µL iTru7 Primer(s) (5 µM -> 0.5 µM final )

10.0 μL clear PCR product from step 12 (placed on magnet).

Cycle: 98°C for 2 minutes.; then, 5-10 cycles of: 98°C for 20 sec., 60°C for 15 sec., 72°C for 30 sec.; followed by 72°C for 5 min. Hold at 15°C. [Note: Adjust the number of cycles based on the amount of input DNA, more cycles if less input DNA was used.]

13. Pool all three reactions from above & purify with SpeedBeads (Thermo-Scientific, Waltham, MA, USA) to a 1:1 ratio, DNA:SpeedBeads. purify as normal and resuspend in 100 µL of TLE.
14. Run 5µL on agarose gel to ensure each sample worked.
15. Quantify with Qubit
