## Supplementary material for "Adapterama II: Universal amplicon sequencing on Illumina platforms (TaggiMatrix)": File S3

**Supplementary methods and results for TaggiMatrix example datasets**

***Kissing Bug Blood-Meal Source Characterization***

In this first project, we characterized the source of blood meals in the kissing bug, *Rhodnius pallescens* from a previous study (Kieran et al. 2017)*.* Kissing bugs were captured from palm trees in Trinidad de las Minas, Capira (8.46713, -79.59451) (N = 10 bugs) and Las Pavas, La Chorrera (9.05448, -79.53352) (N = 7), Panama using Noireau traps (Noireau *et al*. 2002) from June to July 2013 and 2015. We placed bugs in 95% molecular grade ethanol. We dissected guts from the exoskeletons and extracted DNA from the guts using a Qiagen DNeasy Blood and Tissue Kit. We constructed libraries in a three-step library preparation method (Method 1 in Table 3, main text). First, we used standard *cytochrome-b (cyt-b)* primers (i.e., no fusion nor indexes added) to amplify the DNA using PCR conditions described by Parson et al. (2000; Table 2 in main text). We verified PCR products on a 1.5% agarose gel and cleaned them using a 1:1 ratio of SpeedBeads (Thermo Fisher Scientific, Waltham, MA, USA; see Glenn et al*.* 2016 for preparation methods), reconstituting the product in 25 µL TLE. We used 10 µL of this PCR product in a 25 µL KAPA LTP Library Preparation Kit reaction, following the manufacturer’s protocol (KR04353 v2.13; Roche, Basel, Switzerland) with our iTru adapters and primers. We used the iTru universal y-yoke adaptor during the ligation step (Glenn et al*.* 2016). Finally, we conducted a limited-cycle PCR using iTru primers, with each 12.5 µL reaction consisting of 2.5 µL 5x Kapa buffer, 0.375 µL 10 mM dNTPs, 0.25 µL Kapa HiFi polymerase, 1.25 µL i5 (5 µM), 1.25 µL, i7 (5 µM), 1.625 µL H20, and 5 µL of ligated amplicon DNA. Thermocycler conditions were as follows: 98°C for 45 sec, followed by 10 cycles at 98°C for 30 sec, 60°C for 30 sec, 72°C for 30 sec, and a final extension at 72°C for 5 min. We cleaned this product with a 1:1 SpeedBeads:DNA ratio and pooled it with other uniquely-indexed samples prior to sequencing. We sequenced these libraries using an Illumina MiSeq v3 600 cycles kit at the Georgia Genomics Facility. We demultiplexed sequencing data by outer indexes using bcl2fastq (Illumina v1.8.4). We set pairs and merged sequence data using the FLASH v1.2.9 plugin (Magoč & Salzberg, 2011) implemented in Geneious v9.1.2. In the same program, we trimmed adaptors and primers, and filtered for quality using default settings and an error probability limit of 0.01. We then extracted and exported sequences with lengths between 270–280 bp as FASTA files. We created a reference database of *cyt-b* sequences from Panamanian vertebrates using publicly available sequence data, substituting in close relatives of species absent from GenBank. We used MacQIIME v1.9.1 to map sequence reads to this custom database, implementing UCLUST (Edgar, 2010) for picking OTUs and making taxonomic classifications for OTUs with an overall similarity of 97%. Resulting OTU identifications for each sample represented absolute abundance (i.e., total number of reads). Following Kieran et al. (2017), we eliminated species hits receiving ≤10% of the total read hits for each sample for a conservative estimate of blood meal source.

We obtained between 2,276 and 317,515 (mean = 116,902) paired reads for each sample. We identified a total of five unique vertebrate species across all samples with two species comprising 83% of total reads: *Didelphis marsupialis* (65%) and *Mus musculus* (18%) (Tables 1–2). The Las Pavas site had more diversity in blood meal source (5/5 species) compared to the Trinidad de las Minas site (2/5 species). We identified the blood meal source in 16 of the 17 samples. Unassigned reads accounted for 12.5% of the total. Only one sample (TM27) was not identified using the current reference database. Two other samples (Pa24 and Pa72) had many unidentified reads and possibly consist of multiple sources of blood meal.

These three samples account for 95.5% of the unassigned reads, suggesting that this is an accurate and effective method limited by the availability of sequence resources to construct a complete reference database (Kieran et al., 2017).

**Table 1. Mammal OTUs from seven *Rhodnius pallescens* blood-meal samples** **from Las Pavas, Panama**

Values represent raw numbers, and the site total is also shown as a percentage of all reads.

| OTU ID | **Pa112** | **Pa117** | **Pa146** | **Pa15** | **Pa21** | **Pa24** | **Pa72** | **Total** | |
| --- | --- | --- | --- | --- | --- | --- | --- | --- | --- |
| *Diplomys labilis* | 4176 |  |  |  |  |  |  | 4176 | 1.73% |
| *Caluromys derbianus* | 20444 |  |  |  |  |  |  | 20444 | 8.49% |
| *Didelphis marsupialis* |  | 14234 |  |  | 65001 | 37212 |  | 116447 | 48.34% |
| *Coendou bicolor* |  |  |  |  |  |  | 8112 | 8112 | 3.37% |
| *Mus musculus* |  |  | 23507 |  |  |  |  | 23507 | 9.76% |
| Unassigned | 132 | 519 | 8 | 617 | 454 | 11947 | 54526 | 68203 | 28.31% |
| Total | 24752 | 14843 | 23515 | 632 | 69446 | 52924 | 62644 | 240889 | 100% |
| **10% cutoff** | **2475.2** | **1484.3** | **2351.5** | **63.2** | **6944.6** | **5292.4** | **6264.4** |  |  |

**Table 2. Mammal OTUs from seven *Rhodnius pallescens* blood-meal samples** **from Trinidad de las Minas, Panama**

| OTU ID | **TM071** | **TM074** | **TM075** | **TM128** | **TM17** | **TM19** | **TM25** | **TM27** | **TM28** | **TM51** | **Sp. site total** | |
| --- | --- | --- | --- | --- | --- | --- | --- | --- | --- | --- | --- | --- |
| *Didelphis marsupialis* |  |  |  |  | 71138 | 143067 | 43740 |  | 64580 | 41835 | 364,360 | 73.10% |
| *Mus musculus* | 30112 | 27176 | 28264 | 24216 |  |  |  |  |  |  | 109,768 | 22.02% |
| Unassigned | 7 | 3 | 7 | 8 | 118 | 3 | 141 | 21887 | 1489 | 619 | 24,282 | 4.87% |
| Total | 30124 | 27179 | 28405 | 24224 | 71256 | 143072 | 44058 | 22136 | 67280 | 42982 | 498,410 | 100% |
| **10% cutoff** | **3012.4** | **2717.9** | **2840.5** | **2422.4** | **7125.6** | **14307.2** | **4405.8** | **2213.6** | **6728** | **4298.2** |  |  |

Values represent raw numbers, and the site total is also shown as a percentage of all reads.

***Molecular Identification of Pathogenic Fungi on Trees Using ITS***

In this second project, we grew fungal cultures isolated from tree samples collected on Barro Colorado Island, Panama. These cultures consisted of three types: 1) 48 homokaryon cultures grown from single fungal spores sampled from a total of eight tree species; 2) 47 heterokaryon cultures grown from tissue samples of fruiting fungal bodies sampled from a total of 35 tree species; and 3) 48 heterokaryon cultures grown from multi-spore samples collected from a total of 25 tree species. To sample and extract DNA from each culture, we transferred 2–5 pieces of media and fungi to 2.0 mL vials filled with distilled and sterilized water. Under a sterile laminar flow hood, we transferred 2–4 of these pieces and 100 µL of liquid to tubes of 0.1 mm glass beads (Qiagen, Valencia, CA, USA). We then added 400 µL of digestion buffer (10:1 of 1 M Tris-HCL:0.5 M EDTA pH8) to each tube. We disrupted the sample using a FastPrep-24 (MP Biomedicals, China) using three cycles of 60 sec at 4 m/sec. We then centrifuged samples at 10,000 g for one minute. We carefully transferred the supernatant to a sterile 1.5 mL tube and added 25 µL of Proteinase K solution (20 mg/mL). We vortexed samples and incubated them overnight at 56 °C while shaking at 70 rpm. After digestion, we added 800 µL SpeedBeads, cleaned as usual, and rehydrated in 30 µL TLE. We verified the DNA quality by electrophoresis through a 1.25% agarose gel and quantified the DNA extractions with a Qubit Fluorometer or plate reader.

We used TaggiMatrix to generate internally indexed fusion primers to amplify the ITS1 and ITS2 regions. These primers were designed from the following standard primers: ITS1-F_KY02, ITS2_KY02, ITS3_KY02 and ITS4 (Toju et al. 2012; White et al. 1990; Table 2 in main text). We also synthesized “flipped” versions of these tagged fusion primers (Fig. 2). We constructed libraries using Method 2 (Table 3 in main text) with a three-step PCR approach. We created a total of nine 96-well plate pools, each consisting of 47–48 culture samples and two PCR sets corresponding to the different orientation (standard and “flipped”, Fig. 2 in main text). In brief, we conducted a first PCR using ITS1-F_KY02 and ITS4 standard primers, with each 50 μL reaction consisting of: 10 µL 5x Kapa buffer, 1.5 µL 10 mM dNTPs, 1.0 µL Kapa HiFi polymerase, 24.5 µL H20, 4 µL ITS1-F_KY02 primer (5 µM), 4 µL ITS4 primer (5 µM), and 5 µL of the sample DNA. Thermocycler conditions were as follows: 95 °C for 5 min, followed by 10 touchdown cycles of 94°C for 30 sec, 60°C for 30 sec, -1°C/cycle, and 72°C for 30 sec, followed by 10 more cycles of 94°C for 30 sec, 50°C for 30 sec, and 72°C for 30 sec with a final elongation step of 72°C for 5 min.

After this first PCR, we cleaned each product using a 0.8:1 ratio of SpeedBeads:DNA and reconstituted in 50 µL TLE. We then performed six different PCR reactions using indexed fusion primers per sample—three with the standard primer orientation and three with the “flipped” primer orientation. These PCRs used indexed fusion primers derived from the following primer pairs: 1) ITS1-F_KY02 and ITS2_KY02 to amplify the entire ITS1 fragment; 2) ITS3_KY02 and ITS4 to amplify the entire ITS2 fragment; and 3) ITS1-F_KY01 and ITS4 to amplify a partial fragment of ITS1 and ITS2 (Table 2 in main text). Each 25 μL reaction consisted of: 5 µL 5x Kapa buffer, 0.75 µL 10 mM dNTPs, 0.5 µL Kapa HiFi polymerase, 9.75 µL H20, 2 µL forward primer (5 µM), 2 µL reverse primer (5 µM), and 5 µL of the cleaned PCR product. Thermocycler conditions were as follows: 95 °C for 5 min, followed by 10 touchdown cycles of 94°C for 30 sec, 60°C for 30 sec, -1°C/cycle, and 72°C for 30 sec, followed by 5 more cycles of 94°C for 30 sec, 50°C for 30 sec, and 72°C for 30 sec with a final elongation step of 72°C for 5 min. We purified and normalized these products from the second PCR using SequalPrep Normalization Kit (Thermo Fisher Scientific, Waltham, MA, USA) following the manufacturer’s protocol.

Within each plate, we then pooled 10 µL of each eluted product. We dried these pools in a vacuum rotator at 60°C until ~100 µL of pool was left. We then conducted a PCR using iTru primers on each pool. Each 25 μL reaction consisted of: 5 µL 5x Kapa buffer, 0.75 µL 10 mM dNTPs, 0.5 µL Kapa HiFi polymerase, 3.75 µL H20, 2.5µL i5 primer (5 µM), 2.5 µL i7 primer (5 µM), and 10 µL of the pool. Thermocycler conditions were as follows: 95°C for 5 min, followed by 6 cycles of 94°C for 30 sec, 60°C for 30 sec, and 72°C for 30 sec, and a final elongation step of 72°C for 5 min. Each product was verified in 1.5% agarose gel. We cleaned products with a 1:1 SpeedBeads:DNA, quantified them using a Qubit Fluorometer, and pooled them accordingly. We sequenced these libraries using an Illumina MiSeq v3 600-cycle kit at the Georgia Genomics Facility.

We demultiplexed pools by outer tags using Illumina’s bcl2fastq (Illumina v1.8.4). Then, we demultiplexed samples within each pool were demultiplexed by internal tags using Mr Demuxy v1.2.0 (<https://pypi.python.org/pypi/Mr_Demuxy/1.2.0>). We merged paired reads for the first two primer pairs using FLASH2 (Magoč & Salzberg, 2011). For the third primer pair, we analyzed Read1 and Read2 independently. We trimmed primer sequences using BBDuk from the BBMap package v37.22 (<http://sourceforge.net/projects/bbmap/>). We quality-filtered all reads using the fastq_quality_filter pipeline from FASTX_Toolkit v0.0.14 (<http://hannonlab.cshl.edu/fastx_toolkit/>), setting a minimum quality score (*q*) of 30 and a minimum percent of bases that must have *q* quality of 80, and we converted these output files into FASTA files using the tool fastq_to_fasta. We identified and extracted ITS sequences using ITSx v1.0.11 (Bengtsson-Palme *et al*. 2013). We used MacQIIME v1.9.1 to remove chimeras and find OTUs. We used the UNITE fungal dynamic database v7.0 (2016-01-31; Abarenkov et al. 2010) as a reference, and the UCLUST (Edgar, 2010) OTU picking algorithm (pick_open_reference_otus.py), and BLAST (Atschul et al. 2015) for taxonomic assignment. After assigning OTUs for each of the six PCRs in each culture sample, we used the sequencing information from all six PCR designs across each sample (full-ITS1, full-ITS2, incomplete-ITS1 & ITS2, and the same regions using “flipped” primers), and then identified and graphed the OTUs for each sample using Phyloseq v1.19.1 in R v3.3.1 within R Studio v1.0.136.

We obtained between 257,410 and 618,525 (mean = 436,825) reads per plate. After demultiplexing by internal tags, only eight out of the 858 PCRs returned no reads (0.9%), and only four additional PCRs returned fewer than 10 reads (0.5%). Other than these samples, we obtained between 10 and 43,348 (mean = 4,015) reads per PCR per sample. On average, 99.14% of reads were retained after merging, trimming, and quality filtering. For those cases where complete ITS1 or ITS2 fragments were recovered by merging paired reads, an average of 93.95% of the demultiplexed reads were positive for the detection and extraction of the respective ITS fragment. In contrast, in the cases where the whole fragment was amplified but paired reads were not merged, an average of 53.21% of the initial reads contained the expected ITS fragment. In general, Read1 reads were assigned more frequently than Read2 reads. Both the original orientation and “flipped” primers worked successfully. We identified the primary OTU from each culture, but in some cases, more than one OTU was assigned to a culture. This was particularly true in samples where the number of reads per sample were below ~1,000. Also, for a very small subset of samples, there was discordance between taxonomic assignments from ITS1 and ITS2. This ambiguity was later resolved with morphology, but these results are not within the scope of this study.

**Supplemental Figure 1. Stacked bar plot of the relative abundance of fungal OTUs from single-spore homokaryon cultures**

Data are pooled across six PCR designs per sample. Sample numbers correspond to a tree species code and the strain culture number.

**Supplemental Figure 2. Stacked bar plot of the relative abundance of fungal OTUs from fruiting body heterokaryon cultures**

Data are pooled across six PCR designs per sample. Sample numbers correspond to a tree species code and the strain culture number.

**Supplemental Figure 3. Stacked bar plot of the relative abundance of fungal OTUs from multispore heterokaryon cultures**

Data are pooled across six PCR designs per sample. Sample numbers correspond to a tree species code and the strain culture number.

***Plethodontid Salamander Environmental DNA***

In this third project, we developed and tested a protocol to characterize semiaquatic Appalachian salamander communities from environmental DNA (eDNA) samples. We used the program ecoPrimers and mtDNA sequence data from GenBank to design metabarcoding primers to characterize communities of our target taxa (family Plethodontidae)­. Using primers targeting a portion of 12S rRNA gene (Table 2 in main text), we created non-indexed and indexed iTru fusion primers.

To build a new reference database of 12S barcode sequences, we collected one tissue from each of seven species of plethodontid salamanders documented in low-elevation streams at the Coweeta Hydrologic Laboratory in Macon County, North Carolina from a nearby location in Towns County, Georgia. We extracted DNA from tissues using Qiagen DNeasy Blood and Tissue Kits, and we normalized extracted DNA to a concentration of 2 ng/µL. We constructed these libraries using Method 4 (Table 3 in main text). We conducted PCRs using non-indexed iTru fusion primers, with each 25 µL reaction consisting of: 5 µL 5x Kapa buffer, 0.75 µL 10 mM dNTPs, 0.5 µL Kapa HiFi polymerase, 11.75 µL H20, 1 µL forward primer (10 µM), 1 µL reverse primer (10 µM), and 5 µL of the sample DNA. Thermocycler conditions were: 95**°**C for 3 min, 20 cycles of 98**°**C for 20 sec, 60**°**C for 15 sec, and 72**°**C for 30 sec, and then 72**°**C for five min, and held infinitely at 15 **°**C. We cleaned PCR products with SpeedBeads using a 2:1 SpeedBead:DNA ratio, resuspended in 25 µL TLE, and diluted this product 20X in TLE. We then conducted a second, limited cycle PCR using iTru primers, each 25 µL reaction consisting of: 5 µL 5x Kapa buffer, 0.75 µL 10 mM dNTPs, 0.5 µL Kapa HiFi polymerase, 2.5 µL i5 (5 µM), 2.5 µL, i7 (5 µM), 8.75 µL H20, and 5 µL of amplicon DNA. Thermocycler conditions were: 95 **°**C for 3 min, then 16 cycles of 98**°**C for 20 sec, 60**°**C for 15 sec, and 72**°**C for 30 sec, and then 72**°**C for 5 min, and an infinite hold at 15**°**C. We cleaned PCR products again using a 2:1 ratio of SpeedBeads:DNA. We quantified these libraries using a Qubit 2.0 Fluorometer and pooled them for sequencing using an Illumina MiSeq v2 600 cycle kit at the Georgia Genomics Facility. We demultiplexed reads using bcl2fastq (Illumina v1.8.4). We used MacQIIME v1.9.1 to trim and filter reads, and we used AliView 1.18 to generate consensus sequences for each individual.

We collected three 1 L water samples each from three headwater streams at the Coweeta Hydrologic Laboratory in Macon County, North Carolina. We vacuum-filtered each sample and extracted DNA from the filter papers following the protocols described in Pierson et al. (2016). We conducted three separate PCRs for each sample, with each PCR using a different combination of internally indexed iTru fusion primers (Method 5 from Table 3, main text). Otherwise, the recipe and thermocycling conditions for this first PCR were identical to those of the construction of the reference library. After the first PCR, we cleaned PCR products using a 2:1 SpeedBeads to PCR product ratio and resuspended in 25µL TLE. We did not dilute these PCR products before the second PCR. We conducted the second PCR with a recipe and thermocycling conditions identical to those from the construction of the reference library. After the second PCR, we again cleaned PCR products using a 2:1 SpeedBeads:DNA ratio. We then quantified them using a Qubit 2.0 Fluorometer and pooled them. We size-selected for the expected amplicon size on Pippip Prep and pooled these samples with unrelated projected to be sequenced at the Georgia Genomics Facility using an Illumina MiSeq v3 600 cycle kit. We demultiplexed reads using bcl2fastq (Illumina v1.8.4). We used MacQIIME v1.9.1 to trim, filter, and map reads to a reference database created from the seven consensus sequences we generated. We plotted our results using R v3.2.3.

We obtained between 618 and 1,843 (mean = 1,201) reads for each sample used to construct the reference database. We obtained between 28,777 and 323,361 (mean = 163,555) reads for each eDNA PCR replicate. We mapped reads from eDNA samples to 6/7 species in our reference database (Fig. 4), and no reads from any of the negative control replicates mapped to sequences in the reference database.

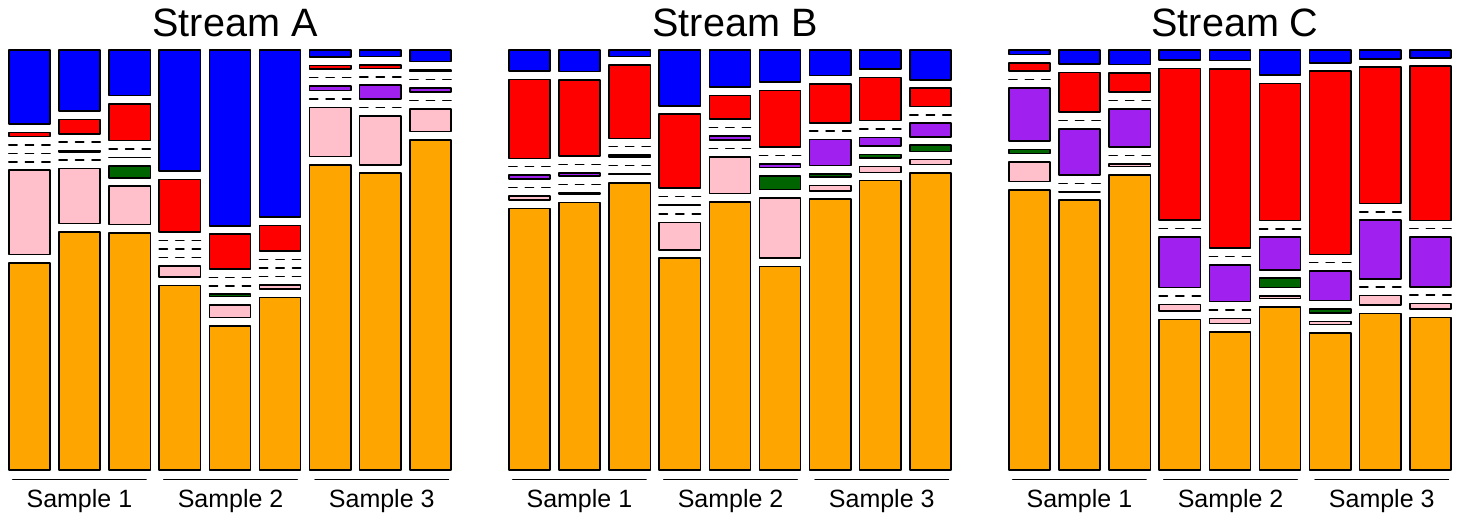

**Figure 4. Stacked bar plot of the relative abundance of reads assigned to six salamander species in eDNA samples**

Data are shown for each of three PCR replicates from each of three samples collected from each of three streams. Species are represented as follows: blue = *Eurycea wilderae*, red = *Gyrinophilus porphyriticus*, purple = *Desmognathus ocoee*, green = *Desmognathus aeneus*, pink = *Desmognathus monticola*, and orange = *Desmognathus quadramaculatus*.

***Differences in methylation of p21 of embryonic kidney cells and freshly isolated human proximal tubule cells***

In this fifth project, we compared basal DNA methylation of *p21* promoter between two different passages of human embryonic kidney (HEK293) cells and two different pools of freshly isolated human proximal tubule (hPT) cells.

We purchased human embryonic kidney cells (HEK293) from the American Type Culture Collection (Manassas, VA). We derived human proximal tubular (hPT) cells from whole, de-identified human kidneys that we obtained through the International Institute for the Advancement of Medicine (Edison, NJ). All tissue was scored by a pathologist as normal (i.e., derived from non-cancerous, non-diseased tissue). We pelleted cells (5x10^6^) at 1000 rpm for 5 min and discarded the supernatant. We extracted genomic DNA using the Qiagen DNeasy Blood and Tissue kit (Valencia, CA) following the manufacturer’s protocol. We eluted DNA two successive steps to obtain a maximum yield, using 120 µL followed by 40 µL of elution buffer. Following quantification using a Nanodrop spectrophotometer, we bisulfite-treated 2 µg of the extracted DNA using the Zymo Research EZ-DNA Methylation Lightning kit (Irvine, CA) following the manufacturer’s protocol, and re-quantified the DNA.

We used this bisulfite converted DNA (350 ng) as template to amplify different regions of the *p21* promoter. We designed and synthesized tagged iTru fusion primers targeting two regions (Table 2). The first locus amplified was a 350 bp fragment of the human p21 promoter region adjacent to the transcription start site (TSS) termed as hp21-TSS. The second locus was a 335 bp fragment including the transcription factor binding site approximately 700 bp upstream of the TSS called the sis-inducible element (SIE-1) termed hp21-SIE1.

We constructed these libraries using Method 4 (Table 3 in main text). We conducted the first PCRs using iTru fusion primers (non-indexed), with each 25 µL reaction consisting of: 2.5 µL of 10x hot start PCR buffer, 3 µL of 25mM MgCl_2_, 0.5 µL of 10 mM of each dNTP, 0.3 µL Maxima hot start Taq DNA polymerase 5U/ µL, 1 µL of 10 µM Forward Primer, 1 µL of 10 µM Reverse Primer, 350ng of DNA template and complete volume with water. The PCR conditions were as follows: initial denaturation at 95°C for 5 min; 40 cycles of 95°C for 30 sec, 61.8°C for 45 sec, and 72°C for 45 sec; final extension at then 72°C for 10 min.

We then separated PCR products by electrophoresis on a 1% (w/v) agarose gel and visualized the products with ethidium bromide under a UV trans-illuminator. We extracted the amplicons corresponding to the loci using Nucleospin gel and PCR clean-up kit (Macherey-Nagel, Bethlehem, PA) following the manufacturer’s instructions. We normalized purified PCR amplicons from the agarose gel extraction to 5 ng/µL. We then performed a limited cycle PCR to attach iTru5 and iTru7 primers with eight nucleotide indexes. The 15 µL limited cycle reaction contained 3 µL of 5X Kapa buffer, 0.45 µL of each dNTP 10 mM, 1.5 µL of each iTru5 and iTru7 primers 3 µM, 0.3 µL of Kapa HiFi HotStart DNA polymerase 1U/µL, and 3 µL of 5 ng/µL of DNA. The reaction conditions were: 98°C for 5 min, 11 cycles of 98°C for 15 sec, 60°C for 30 sec and 72°C for 30 sec and a final 72°C for 1 min. We purified these products using 1:1 SpeedBeads:DNA ratio and pooled them with libraries from unrelated projects for sequencing on an Illumina MiSeq v3 600-cycle kit at the Georgia Genomics Facility.

We demultiplexed paired-end reads using the Illumina software bcl2fastq (Illumina v1.8.4). We conducted methylation analysis using Bismark (Krueger et al*.*, 2011) in three steps . First, we indexed the reference sequence available from NCBI (Gene ID: U24170.1) using Bowtie 2 (Langmead & Salzberg, 2012). Second, we mapped and aligned only Read 1 reads against the reference, producing a BAM file. And third, we extracted the methylation profile from the BAM file. Finally, we calculated the CpG site specific methylation percentage.

We obtained an average of approximately 10,000 reads per sample and compared the basal DNA methylation of *p21* promoter in HEK293 cells and hPT cells and observed similarities in the region adjacent to the transcription stat site and differences in the sis-inducible element (SIE-1), a transcription factor binding site. The average methylation of hp21-TSS in hPT cells was 1.4% and not significantly different from that in HEK293 cells (Fig. 5a). In contrast, the average methylation of all three CpG sties in hPT cells were lower than that measured in HEK293 cells at the hp21-SIE1 site (Fig. 5b).

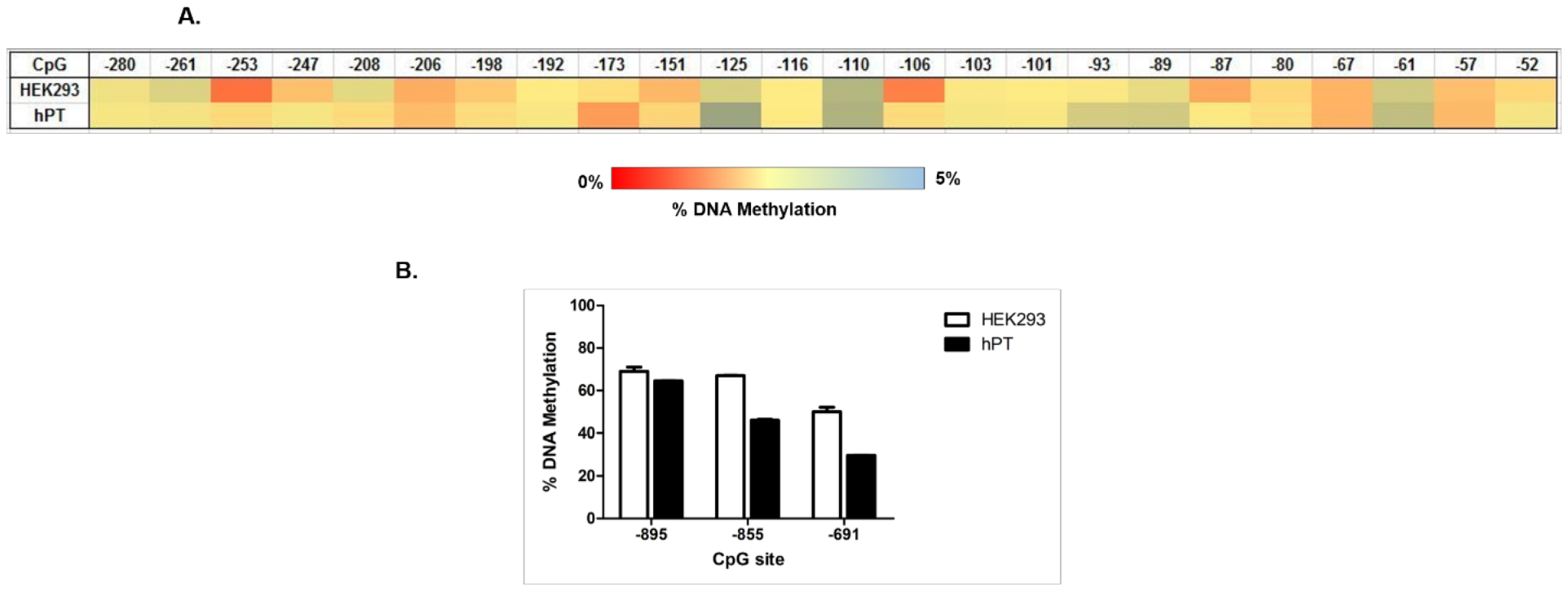

**Figure 5.** **Comparison of basal DNA methylation of the p21 promoter region between HEK293 cells and freshly isolated human proximal tubule (hPT) cells**

(A) Heat-map of the site-specific percent DNA methylation changes as determined by TGBS in the human p21 promoter region at the transcription start site (hp21-TSS). Heat map intensity is showed in the sidebar with deep red indicating percent methylation value towards zero and pale blue indicating relatively higher methylation of 5%. (B) Comparison of methylation of human p21 promoter at the transcription factor binding site SIE-1 between HEK293 and hPT cells. Data are represented as the mean ± SEM of two different passages of HEK293 cells and two different pools of the hPT isolated cells (n=2).

***Bacterial Gut Microbiome from Cotton Mice***

In this fourth project, we characterized bacterial communities from the intestines of 90 wild cotton mice (*Peromyscus gossypinus*) to examine the influence of chronic heavy metal exposure. We selected a universal primer pair targeting a fragment size of approximately 444 bp in the 16S rRNA gene, a common barcoding locus for bacteria (Klindworth et al. 2013; Table 2 in main text). We used TaggiMatrix to design and synthesize iTru fusion primers. We extracted DNA from 50 fecal pellets collected during necropsy using a PowerSoil DNA Isolation Kit (MO BIO Laboratories, Carlsbad, CA). We quantified the extracted DNA using a Qubit 2.0 fluorometer and stored it at -80°C.

We constructed these libraries using Method 5 (Table 3 in main text). We conducted the first PCRs using internally indexed iTru fusion primers, with each 25 µL reaction consisting of: 5 µL 5x KAPA HiFi buffer, 0.75 µL 10 mM dNTPs, 0.5 µL KAPA HiFi HotStart, 1.5 µL 5µM Forward Primer, 1.5 µL 5 µM Reverse Primer, 13.75 µL H20, 2 µL of microbial DNA. The PCR conditions were as follows: initial denaturation at 95°C for 3 min; 20–25 cycles of 95°C for 20 sec, 60°C for 30 sec, and 72°C for 30 sec; final extension at then 72°C for 5 min.

We purified PCR products, quantified them using a Qubit fluorometer, and diluted them to a final concentration of 2 ng/μL using sterile, nuclease-free water. We used the diluted PCR amplicons in a second PCR reaction containing the TruSeq Illumina P5 and P7 primers. The PCR amplification conditions were as described above, but for only fourteen cycles. We purified these quadruple-indexed libraries using a 2:1 SpeedBeads:DNA ratio and pooled them with other libraries from unrelated projects. We sequenced these libraries at the Georgia Genomics Facility using an Illumina MiSeq v3 600-cycle kit.

We initially demultiplexed reads using bcl2fastq (Illumina v1.8.4), then used Mr_Demuxy (https://pypi.python.org/pypi/Mr_Demuxy/1.2.0) to demultiplex combinatorially tagged samples. We used Geneious v10.0 (Biomatters, Auckland, New Zealand) to quality-trim and set paired-reads with an expected insert size of 444 bp and merged using the FLASH v1.2.9 plugin (Magoc & Salzberg, 2011). We then select edreads between 300-400 bp and removed low-quality reads using FASTX_Toolkit v0.0.14 (http://hannonlab.cshl.edu/fastx_toolkit/).

We used MacQIIME v1.9.1 to map reads to the Greengenes v13_8 16S reference database ([http://greengenes.secondgenome.com/](https://www.dropbox.com/referrer_cleansing_redirect?hmac=JIbY%2FOd9lIoYkXfvvc93xM%2B1cP6IKk7LoDnYEROU5ks%3D&url=http%3A%2F%2Fgreengenes.secondgenome.com%2F); DeSantis et al. 2006) using the UCLUST (Edgar, 2010) OTU picking algorithm and the default settings of ≥97% similarity to reference sequences for taxonomy assignment. We estimated community composition, alpha-, and beta-diversity using Phyloseq v1.19.1 in R v3.3.1 and Primer-E v7.

After completing all quality filtering steps, de-noising, and chimera removal, we compiled 7,712,212 high quality (Q≥ 20)16S rRNA gene sequences with an average of 412.7 ± 29.2 reads per sample. We conducted analyses on rarefied data using an even sampling depth of 15,000 reads per sample. We recovered a total of 90,862 OTUs (3,827 ± 2,035 OTUs/sample) spanning 10 bacterial phyla commonly associated with the mammalian gastrointestinal (GI) tract.

Generally, gut microbiome samples from sites with high levels heavy metal concentrations had the lowest observed bacterial diversity, while those from sites with the lowest heavy metal concentrations had the highest observed bacterial diversity. This was true across a wide range of diversity metrics (Fig. 6). Pairwise group comparisons revealed significant differences in Chao1 and Faith’s phylogenetic diversity between gut microbiome communities from sites high and low concentrations of heavy metals (non-parametric test, P < 0.05).

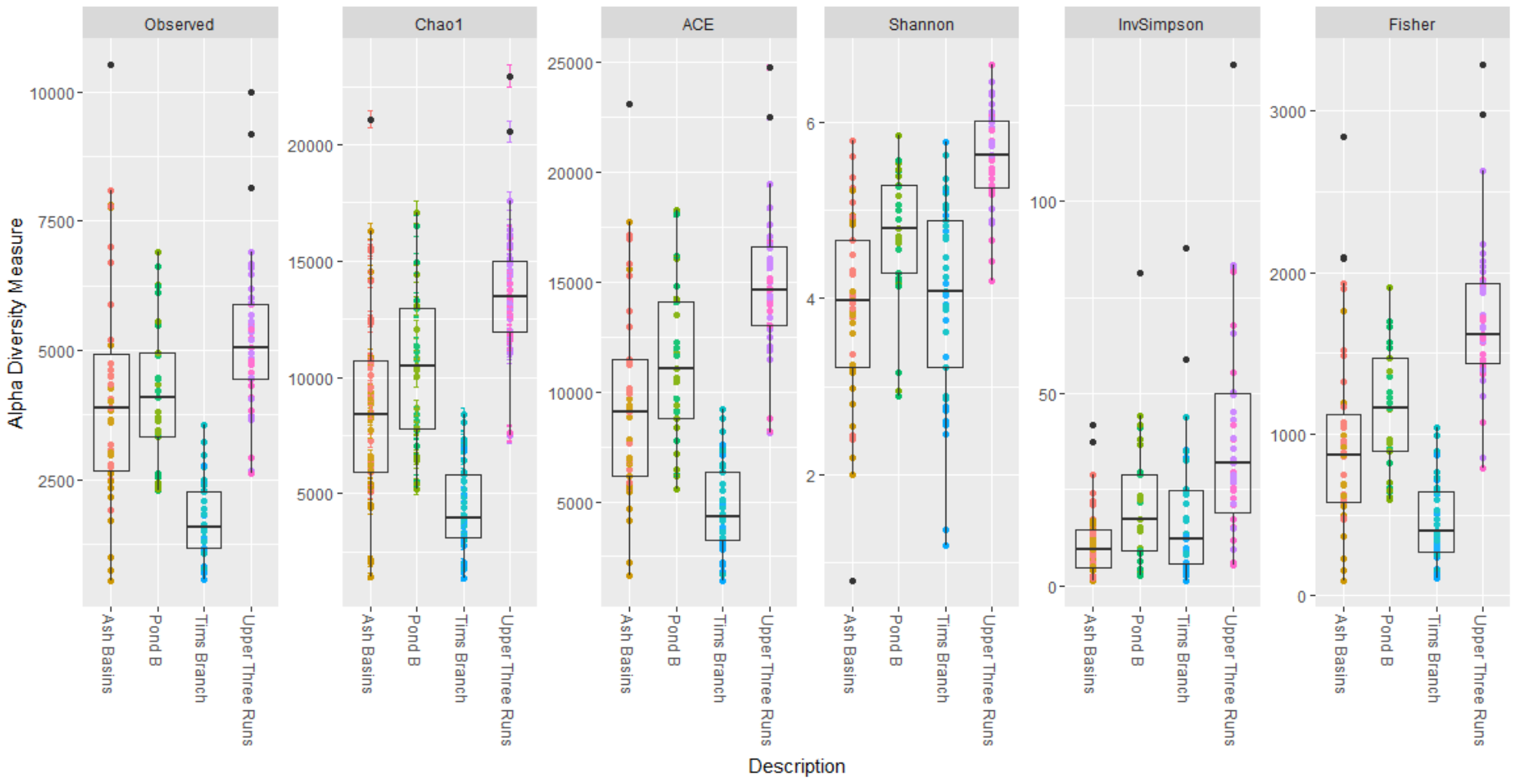

**Figure 6. Observed bacterial OTUs and bacterial alpha diversity estimates for four sampling sites with high and low content of heavy metals**

Ash Basin and Tim’s Branch are sites with high concentrations of heavy metals. Pond B and Upper Three Runs are sites with low levels of heavy metals. Number of bacterial OTUs observed, Chao1, ACE, Shannon, Inverse Simpson, and Fisher diversity measures are presented.

**Wisteria *population genetics***

In this sixth project, we selected five primer pairs targeting one chloroplast locus, two mitochondrial loci, and two nuclear loci for fine-scale population genetic analysis of an invasive vine in the genus *Wisteria* (Trusty *et al.* 2007a, 2007b, 2008). We synthesized a total of 10 indexed Nextera fusion primers. We prepared libraries from 95 individuals distributed across 25 populations along two major roads in Athens, Georgia, USA. For the first PCR, we prepared a single 12.75 µL PCR reaction per locus per individual *Wisteria*. Each PCR reaction mix included 2.5 µL KAPA HiFi Fidelity Buffer (Roche, Basel, Switzerland), 0.38 µL of 10 mM dNTPs, 1.4 µL for both the forward and reverse Nextera indexed fusion primers at 10 µM, 0.25 µL of KAPA HiFi HotStart DNA polymerase, and 2.5 µL of 10 ng/uL DNA. The PCR conditions were as follows: initial denaturation at 95°C for 3 min; 30 cycles of 98°C for 20 sec, 64°C for 15 sec, and 72°C for 15 sec; final extension at 72°C for 5 min. We normalized PCR products among individuals and within each locus based on the brightness of each band on an agarose gel. We performed a second 12.75 µL PCR for each individual and each locus to add iNext5 and iNext7 primers (Glenn *et al*. 2016) to amplicons, with 2.5 µL KAPA HiFi Fidelity Buffer, 0.38 µL of 10 mM dNTPs, 0.4 µL for both the forward and reverse primers at 10 µM, 0.25 µL of KAPA HiFi HotStart DNA polymerase, and 1.0 µL of the previous cleaned-PCR amplicon. This PCR was run for 95°C for 3 min; 10 cycles of 98°C for 20 sec, 60°C for 15 sec, and 72°C for 15 sec; final extension at 72°C for 5 min. We used a SequalPrep Normalization Kit (Thermo Fisher Scientific, Waltham, MA, USA) to normalize and purify the PCR products. We then pooled, quantified, and sequenced final libraries on an Illumina MiSeq v2 500 cycle kit at the Georgia Genomics Facility.

We mapped reads to a reference sequence (generated from Sanger sequencing of 16 samples across eight populations) in Geneious v6 (Biomatters, Auckland, New Zealand). We created a *de novo* assembly to serve as a reference contig for each locus. We identified sequence variation in individuals in Geneious using a minimum coverage of 5 and minimum variant frequency of 0.05. Diversity among samples along each road was analyzed in GenoDive v2.0b23 (Meirmans & Van Tienderen, 2004).

The amplicon size of each locus ranged from 440 to 829 (Table 3), including ~140 bp of Illumina adapter sequences and tags. We recovered a total of 161,228 read pairs of 300 nt, 97.8% of which were mapped to reference sequences (Table 3). From the 95 samples, we recovered an average of 1,697 reads (range: 372 to 4,388) per individual. The nuclear region 997 had 16 variable sites, including the SNP that alters the HpyCH4 IV restriction enzyme recognition site used to distinguish *W. floribunda* from *W. sinensis*. At this site, 5.6% of reads from 2 individuals contained *W. floribunda* alleles. The *cyt-b* sequence had 6 variable sites, including the species-specific AseI restriction enzyme recognition site. Variation in this site indicated that 7.6% of reads from 6 individuals has *W. floribunda* ancestry. Two of these individuals had the same multilocus genotype, consistent with wisteria’s ability to reproduce vegetatively. When distinguishable, the majority of samples contained *W. sinensis* alleles. The nuclear region 824 had 20 SNPs. The plastid sequence had 12 variable sites and the species-specific deletion in *W. sinensis* could not be positively identified due to relatively short read length. The *NAD4* mitochondrial sequence contained 2 variable sites, neither of which created the *W. floribunda* specific SalI restriction enzyme recognition site. The average heterozygosity for all loci was 0.014; the average G_ST_ for all loci was 0.392; and the average G_IS_ was 0.915. Repeated multilocus genotypes were not restricted to the same population and spatial genetic structure was not detected.

**Table 3. Summary data of targeted loci in Wisteria samples**

Amplicon length, percent of reads mapped to the reference, and number of variable sites for each of the 5 loci sequenced in 95 *Wisteria* individuals

| **Locus** | **Amplicon length** | **% reads mapped to reference** | **Number of variable sites** |
| --- | --- | --- | --- |
| nuclear region 824 | 829 | 2.1 | 20 |
| nuclear region 997 | 590 | 17.3 | 16 |
| *trnL* and spacer | 440 | 32.4 | 12 |
| *cyt-b* | 588 | 27.3 | 6 |
| *NAD4* | 575 | 18.7 | 2 |
