## Supplementary material for "Adapterama II: Universal amplicon sequencing on Illumina platforms (TaggiMatrix)": File S5

### Demultiplexing Internal Indexes Using Mr. Demuxy

Author: BadDNA Lab, UGA

Lates version of Mr. Demuxy is available at: https://pypi.org/project/Mr_Demuxy/

Mr. Demuxy was developed by Daniel E. Lefever

#### Goal

If TaggiMatrix from *Adapterama II* system with indexed fusion primers was used to construct libraries, Mr. Demuxy can be used to quickly and efficiently demultiplex samples by internal indexes from Read 1 and Read 2 fastq files.

#### Files to Run Mr. Demuxy

##### 1. Mr. Demuxy uncompressed folder

##### 2. Read 1 and Read 2 fastq Files, uncompressed

##### 3. Indexes txt Files (forward and reverse)

Each set of indexes (Read 1 and Read 2) can be saved as .txt file

**Read 1 Indexes (See Supplemental File S1)**

A GGTAC
B CAACAC
C ATCGGTT
D TCGGTCAA
E AAGCG
F GCCACA
G CTGGATG
H TGATTGAC

**Read 2 Indexes (See Supplemental File S1)**

1 AGGAA
2 GAGTGG
3 CCACGTC
4 TTCTCAGC
5 CTAGG
6 TGCTTA
7 GCGAAGT
8 AATCCTAT
9 ATCTG
10 GAGACT
11 CGATTCC
12 TCTCAATC

#### How to Run Mr. Demuxy

```
python pe_demuxer.py -r1 PATH-TO-READ-1-FASTQ-FILE.fastq \
-r2 PATH-TO-READ-2-FASTQ-FILE.fastq -r1_bc Read_1_indexes.txt \
-r2_bc Read_2_indexes.txt -o PATH-TO-OUTPUT-FOLDER/
```

#### Output of Mr. Demuxy

You will have in the output folder Read 1 and Read 2 fastq files, which names are going to correspond to the combination of internal indexes demultiplexed. For example:

A1_R1.fastq A1_R2.fastq (For indexes A and 1)
A2_R1.fastq A2_R2.fastq (For indexes A and 2)
A3_R1.fastq A3_R2.fastq (For indexes A and 3)
... ...
... ...
H10_R1.fastq H10_R2.fastq (For indexes H and 10)
H11_R1.fastq H11_R2.fastq (For indexes H and 11)
H12_R1.fastq H12_R2.fastq (For indexes H and 12)

#### If changing the names of files is desired:

##### *How to rename output with meaningful names*

Similarly to the last part of the demultiplexing tutorial of external indexes from Brant Faircloth Lab (https://faircloth-lab-documentation.readthedocs.io/en/latest/protocols-computer/sequencing/sequencing-demultiplex-a-run.html). You will need to create a tab-delimited file thas has the index combination name (i.e A1, A2, ..., H11, H12) in column 1 and the names you want for the file in column 2.

For example (temp-names.txt):

A1 Sample_W
A2 Sample_X
...
...
H11 Sample_Y
H12 Sample_Z

Then, you can rename files by using:

```
while IFS=$'\t' read -r column1 column2; do
mv ${column1}_R1.fastq ${column2}_${column1}_R1.fastq;
mv ${column1}_R2.fastq ${column2}_${column1}_R2.fastq;
done < "temp-names.txt"
```
